# Deciphering the ‘m^6^A code’ via quantitative profiling of m^6^A at single-nucleotide resolution

**DOI:** 10.1101/571679

**Authors:** Miguel Angel Garcia-Campos, Sarit Edelheit, Ursula Toth, Ran Shachar, Ronit Nir, Lior Lasman, Alexander Brandis, Jacob H. Hanna, Walter Rossmanith, Schraga Schwartz

## Abstract

N6-methyladenosine (m^6^A) is the most abundant modification on mRNA, and is implicated in critical roles in development, physiology and disease. The ability to map m^6^A using immunoprecipitation-based approaches has played a critical role in dissecting m^6^A functions and mechanisms of action. Yet, these approaches are of limited specificity, unknown sensitivity, and unable to quantify m6A stoichiometry. These limitations have severely hampered our ability to unravel the factors determining where m6A will be deposited, to which levels (the ‘m6A code’), and to quantitatively profile m6A dynamics across biological systems. Here, we used the RNase MazF, which cleaves specifically at unmethylated RNA sites, to develop MASTER-seq for systematic quantitative profiling of m6A sites at 16-25% of all m6A sites at single nucleotide resolution. We established MASTER-seq for orthogonal validation and *de novo* detection of m6A sites, and for tracking of m6A dynamics in yeast gametogenesis and in early mammalian differentiation. We discover that antibody-based approaches severely underestimate the number of m6A sites, and that both the presence of m6A and its stoichiometry are ‘hard-coded’ via a simple and predictable code within the extended sequence composition at the methylation sites. This code accounts for ~50% of the variability in methylation levels across sites, allows excellent *de novo* prediction of methylation sites, and predicts methylation acquisition and loss across evolution. We anticipate that MASTER-seq will pave the path towards a more quantitative investigation of m6A biogenesis and regulation in a wide variety of systems, including diverse cell types, stimuli, subcellular components, and disease states.

## Introduction

M6A is the most abundant modification on mRNA. Although discovered nearly five decades ago, the inability to map m6A on mRNA imposed strong limitations for functionally dissecting its roles. Six years ago, we and others developed immunoprecipitation-based approaches coupled with high-throughput sequencing (m6A-seq, m6A-MeRIP), allowing to detect regions harboring m6A (‘m6A peaks’) (Dominissini et al., 2012; Meyer et al., 2012). These approaches paved the way to major advances in the understanding of m6A, its distribution and conservation, and have facilitated the functional and mechanistic dissection of m6A in development and disease (reviewed in (Knuckles and Bühler, 2018; Meyer and Jaffrey, 2017; Schwartz, 2016; Yue et al., 2015)).

Although antibody-based approaches were pivotal in extending our understanding of m6A, they have several important limitations. First, the *specificity* of detection of m6A sites is limited; In yeast ~50% of the sites that are strongly enriched upon immunoprecipitation result from antibody promiscuity (Schwartz et al., 2013). The anti-m6A antibody also reacts with related modifications, such as m6Am (Dominissini et al., 2012; Linder et al., 2015; Schwartz et al., 2014). Moreover, the *sensitivity* of m6A detection using antibodies could thus far not be evaluated, in the absence of an *orthogonal* technique allowing independent systematic profiling of m6A. Thus, there is a critical need for antibody-independent methods for detection. Second, antibody-based approaches are of limited utility for *quantification* of m6A stoichiometry, i.e. to assess the fraction of modified transcripts. The ability to quantify m6A stoichiometry is critical for functional prioritization of m6A sites and for addressing critical questions pertaining to the biogenesis, regulation, and dynamics of m6A within cells and across stimuli (Grozhik and Jaffrey, 2018; Meyer and Jaffrey, 2014; Schwartz, 2016). Third, the classical m6A-seq approaches are of limited *resolution*, as they provide a sequence window ranging from 3 (Schwartz et al., 2013) to dozens of nucleotides (Dominissini et al., 2012; Meyer et al., 2012) in which m6A is likely to be present. Variants of m6A-seq have been developed, relying on crosslinking of the anti-m6A antibody to the RNA, which upon reverse transcription lead to mutations and truncations at the vicinity of the modified base, achieving nearly single base resolution (Ke et al., 2015; Linder et al., 2015). Nonetheless, the resultant patterns of misincorporation and truncation are complex, diffuse over a 3-4 bp window, and can be variable from one site to another (Ke et al., 2015; Linder et al., 2015). Finally, given the need to immunoprecipitate the RNA, the *starting amounts* of mRNA required for typical m6A-seq libraries - and even more so for cross-linking based derivatives - are limiting. Typically these approaches require micrograms of polyadenylated starting material (Dominissini et al., 2012; Ke et al., 2015; Linder et al., 2015; Meyer et al., 2012; Schwartz et al., 2013), which prohibited interrogation of m6A levels in purified populations of cells, clinical settings, or within specific subcellular compartments. While optimized protocols substantially reduced the starting amounts of mRNA (Merkurjev et al., 2018; Zeng et al., 2018), no protocol exists providing single nucleotide resolution mapping from limited starting mRNA material.

A key, currently poorly understood question pertains to the specificity of the presence of m6A (Darnell et al., 2018; Meyer and Jaffrey, 2017), to which we refer as the ‘m6A code’. While m6A is widespread, its deposition appears to be highly selective, as the sequence motif at which m6A is present, in mammals often represented as DRACH (D=A/G/U, R=A/G, H=A/C/U) is far more abundant than the currently identified m6A sites. DRACH motif is expected by chance roughly every 50-60 nt, amounting to ~45 m6A sites in an average mammalian transcript of 2.5 kb. However, antibody-based approaches detect ~20 fold fewer m6A sites in a typical transcript (Dominissini et al., 2012; Meyer et al., 2012). Moreover, even when present, the stoichiometry of m6A at distinct sites likely varies quite substantially, as could be inferred from careful quantification of m6A levels at a limited number of sites (Horowitz et al., 1984; Liu et al., 2013). Why is m6A present at certain DRACH sites but not at others? And what determines the levels of m6A at these sites? Conceptually, different solutions can be envisioned for this problem of ‘missing specificity’: One scenario is that in addition to a DRACH element acting ‘in cis’, there is substantial ‘trans’ modulation of m6A levels at multiple levels, including its deposition, decoding, and removal (Fu et al., 2014; Wang et al., 2014; Zhao et al., 2018). Under this scenario, the seemingly selective presence of m6A reflects the tight site-specific regulation to which it is subjected. A second scenario is that the ‘missing specificity’ merely reflects our inability to accurately and quantitatively probe m6A distribution and our incomplete knowledge regarding the precise motif recognized by the enzymatic machinery installing m6A. Under this second scenario m6A levels may be *hard-coded* into the mRNA sequence, i.e., the mRNA sequence will dictate the *presence* and the *level* of methylation. Distinguishing between these two scenarios is of critical importance for understanding the key determinants underlying m6A levels, the potential for dynamic modulation of m6A, and the constraints on the evolution of new m6A sites. The inability to quantitatively measure m6A levels at a broad scale has precluded a systematic dissection and modeling of the m6A code.

Here, we develop **M**6**A**-**S**ensi**T**iv**E R**NA digestion and sequencing, MASTER-seq, for antibody-independent detection and quantification of m6A in a systematic scale and at single nucleotide resolution. MASTER-seq builds on the ability of the MazF RNase to cleave RNA at unmethylated sites occurring at ACA motifs, but not at their methylated counterparts (Imanishi et al., 2017). While this approach does not allow interrogation of *all* m6A sites, it allows quantitatively interrogating 16-25% of all methylation sites, and at single nucleotide resolution. We establish the ability of MASTER-seq for *de novo* detection of m6A, for calibrating the sensitivity and specificity of antibody-based approaches for monitoring m6A, and for tracking of m6A dynamics in yeast and mammalian systems. We further reveal that deposition of m6A in yeast is highly deterministic: Both the deposition and the levels (stoichiometry) of m6A are encoded via a simple, predictable sequence composition, which we extensively validate. Changes in sequence composition, as occur naturally throughout evolution, lead to predictable changes in methylation levels, across thousands of sites. The set of orthogonally validated m6A sites in yeast and in mouse, along with the stoichiometries estimated for them, will serve as a critical high confidence resource for the community, and as a reference point for future technologies aiming to quantify m6A levels. We anticipate that MASTER-seq will pave the path towards quantitative investigation of m6A regulation in a wide variety of additional systems, including diverse cell types, stimulations, subcellular compartments, and disease states.

## Results

To obtain quantitative readouts on m6A levels at single nucleotide resolution in an antibody-independent manner, we developed MASTER-seq (Fig. 1A). The approach relies on the ability of the bacterial RNase MazF to cleave RNA immediately upstream of an ‘ACA’ sequence, but not upstream of ‘m6A-CA’ (Imanishi et al., 2017). We reasoned that the RNA cleavage patterns following MazF based cleavage would allow inferring the methylation status at individual residues. Our experimental approach includes the following key steps: (1) Digestion of mRNA with MazF; (2) end repair and ligation of an adapter to the 3’ of the resultant RNA fragments; (3) Reverse transcription, primed from the ligated adapter; (4) Ligation of a second adapter to the 3’ of the cDNA; (5) cDNA Amplification by PCR followed by paired-end sequencing (Fig. 1A, **top**). In an idealized scenario, following MazF treatment, each fragment should begin with an ‘ACA’ site (5’ ACA), and terminate immediately prior to a downstream ‘ACA’ site (3’ ACA). Thus, each pair of sequencing reads - which together capture the precise start and end of the original RNA fragment - provides an indication that two ACA sites (at the 5’ and 3’ of the interval spanning the read pair) were unmethylated in one particular molecule. An m6A containing site is anticipated to be characterized by an abundance of reads passing through - but not terminating - at it (Fig. 1A, **bottom)**. To identify and quantify such methylated sites, we developed MASTER-MINE, a computational pipeline that quantifies the number of reads that begin, terminate, and read-through each transcriptomic ‘ACA’ site. Each site is assigned a 5’ and a 3’ cleavage efficiency, quantifying the number of reads beginning at, or ending immediately before, each ‘ACA’ site, divided by the number of reads overlapping the site respectively (Fig. 1A, **bottom**). In addition, an aggregated cleavage efficiency (‘cleavage efficiency’) is calculated as a mean of the 5’ and 3’ cleavage efficiencies, weighted by the number of reads contributing towards each measurement (‘Methods’).

**Figure 1.**
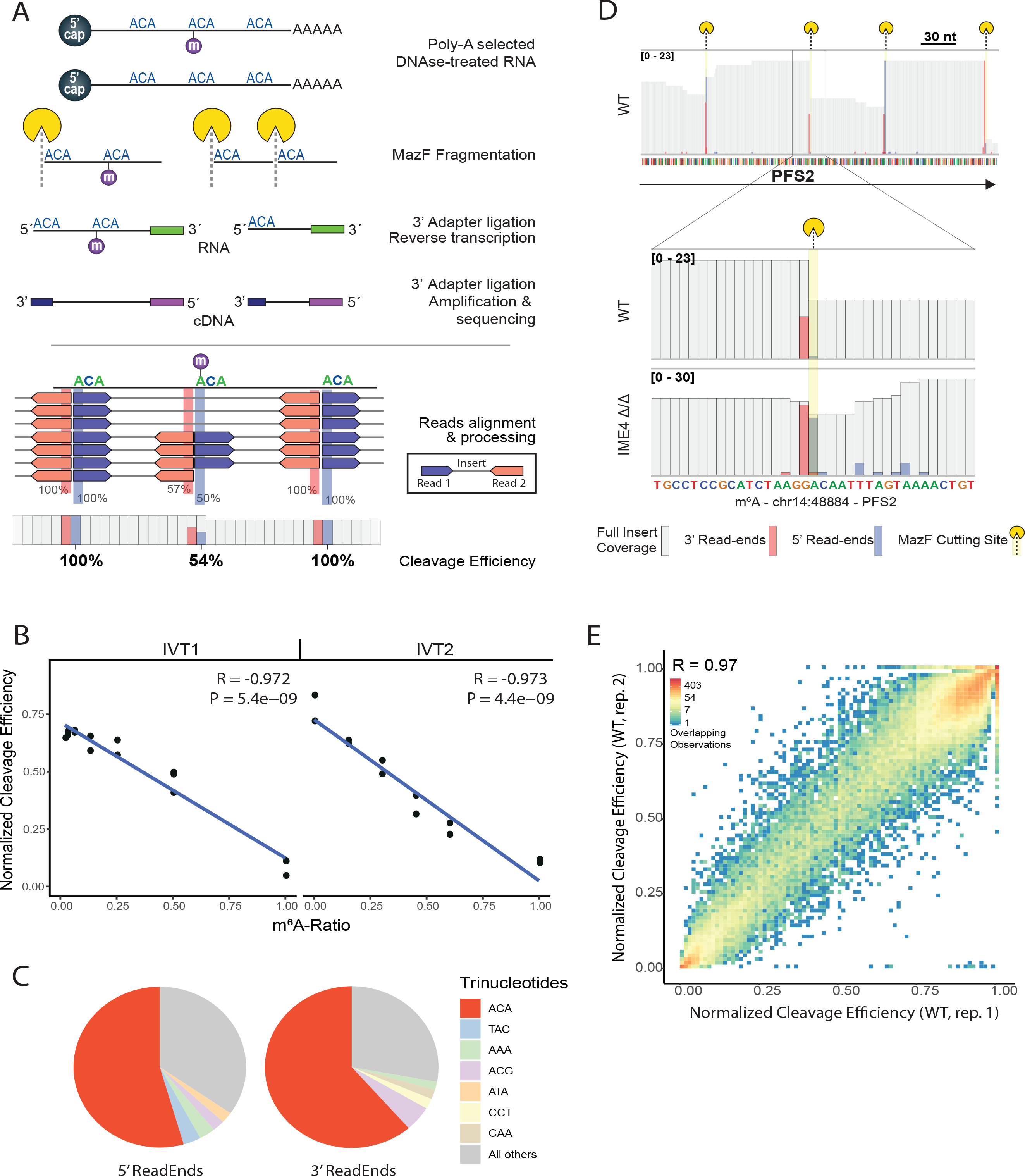
Establishing MASTER-seq. **(A)** Experimental (top): mRNA is Poly-A selected and DNAse treated, followed by MazF digestion, which cleaves RNA at ACA sequences. Ends are repaired and an adapter is ligated to the 3’-end, which enables reverse transcription, and subsequent ligation of a second adapter to the resulting cDNA 3’-end. Finally, amplification and paired-end strand-specific sequencing are performed. Computational (bottom): Reads are aligned to the reference genome. 3’-read-ends and 5’-read-ends are retrieved for relevant coordinates as well as single-base coverage. Cleavage efficiency is calculated by the ratio between the reads starting at an ACA sequence or ending one base downstream it, and their respective single-base coverages. **(B)** MASTER-seq has quantitative qualities. Using synthetic RNAs with a single ACA sequence, either methylated or not, we generated gradients of stoichiometries and spiked them into more complex samples. MASTER-seq measurements agreed with the generated gradients and fit a linear regression with high agreement, denoted by the high R^2^s calculated. **(C)** Relative frequency of the most frequent trinucleotides at the 5’ and 3’ termini of aligned reads. **(D)** Alignment patterns at a methylated site (previously identified based on m6A-seq and *de novo* using MASTER-seq) presented both in WT yeast cells and IME4Δ/Δ cells, both under prophase conditions. The number of sequence tags beginning, ending and overlapping each site are depicted in blue, red, and white, respectively. The mazF cutting site is highlighted in yellow. **(E)** Cutting efficiencies between replicates are highly reproducible. Paired measurements of normalized cleavage efficiency are mapped in each axis to show the level of agreement between replicates. The color gradient ranging from red-yellow-blue depicts the density of overlapping points.

MASTER-seq is thus limited to quantifying m6A sites occurring at ACA motifs. All methylated adenosines are invariably followed by a C in both human and yeast (Dominissini et al., 2012; Harper et al., 1990; Horowitz et al., 1984; Meyer et al., 2012; Wei and Moss, 1977). In yeast, the most prevalent nucleotide at position +2 with respect to the modified position is an ‘A’, present in ~50% of all methylation sites (Schwartz et al., 2013). In mammalian systems ‘A’ is the second-most prevalent nucleotide at this position, present at roughly one-third of the detected m6A sites (Linder et al., 2015). MASTER-seq is thus applicable to a considerable subset of methylation sites, but does not allow *global* mapping/quantification of m6A.

To assess the potential of MASTER-seq to quantitatively capture methylation levels, we applied MASTER-seq to two synthetic short (88 nt long) RNA molecules harboring a single methylation site within a MazF consensus sequence (a single ACA or m6A-CA sequence), which were spiked into complex RNA samples at varying stoichiometries. We obtained excellent agreement between the generated m6A stoichiometries and the experimentally derived cleavage efficiencies (R= −0.97) (Fig. 1B), demonstrating the quantitative power of MASTER-seq under idealized and controlled settings.

### MASTER-seq allows detecting and quantifying m6A levels at endogenous sites

We next tested the ability of MASTER-seq to assay m6A levels at endogenous sites in yeast undergoing meiosis. Yeast mRNAs lack m6A under vegetative growth conditions. In meiosis, the m6A methyltransferase Ime4 induces a widespread methylation program, peaking at prophase (Agarwala et al., 2012; Clancy et al., 2002)(Schwartz et al., 2013). To synchronize meiosis, we applied MASTER-seq to an ndt80Δ/Δ yeast strain, which is genetically synchronized at meiotic prophase, as Ndt80 is required for entry into the meiotic divisions (Brar et al., 2012; Schwartz et al., 2013). Consistent with our expectation, 50-60% of sequenced reads began at ‘ACA’ sites and a similar percentage terminated immediately prior to ACA sites (Fig. 1C). Mapping of the sequencing reads therefore resulted in sharp pile-ups of reads beginning at an ACA sequence and terminating immediately before the next ACA sequence (Fig. 1D). We next examined the agreement between 5’ and 3’ cleavage efficiencies, with the expectation that in an idealized scenario these two measurements - for a single site - should yield an identical value. We found that accurate quantification of 5’ and 3’ cleavage efficiencies is dependent, in a highly predictable manner, on the distance between the interrogated site and its nearest downstream and upstream ACA sites. This is anticipated: if an ‘ACA’ site is within too close proximity of a downstream ‘ACA’ site, it will limit (or prohibit) the ability to capture and sequence accurately the short fragment spanning the two ACAs, and as such the 5’ fragmentation score will yield an underestimate of the cleavage efficiency. Conversely, if an ACA site is preceded by a too close upstream ACA, the 3’ cleavage score will be underestimated **(Fig. S1A-B)**. Accordingly, for the calculation of the cleavage efficiency, MASTER-MINE first defines a set of ‘quantifiable’ ACA sites, i.e. the subset of ACA sites that are within appropriate distances from either an upstream or downstream neighboring ACA sites allowing accurate quantitation, and ‘cleavage efficiency’ scores are calculated only for this subset of sites (Methods). In yeast, 113,014 out of 226,058 ACA sites (50%) are considered quantifiable by MASTER-MINE. The requirement for appropriately distanced ACA sites, along with the above requirement for an ‘ACA’ motif (also present at 50% of the m6A sites) thus renders ~25% of all m6A sites in yeast (and ~16% in mammals) amenable to quantification via MASTER-MINE. Finally, we observed that the cleavage efficiency scores were highly reproducible across biological replicates (R=0.97, Fig. 1E).

We next applied MASTER-seq to mRNA originating from WT and ime4Δ/Δ strains, both in the background of the ndt80Δ/Δ deletion; Deletion of ime4 results in complete elimination of m6A (Schwartz et al., 2013). MASTER-seq was either applied directly to the mRNA (‘Input’) or - as an additional control - to the same mRNA after subjecting it to an m6A-IP, in order to pre-enrich for m6A-containing mRNA. Of note, the IP step is not required for *quantification* of m6A stoichiometry at predefined sites, a key utilization of MASTER-seq. However, applying MASTER-seq to mRNA that had been subjected to m6A-IP step is beneficial for the purpose of *de novo* detection of m6A sites and for QCing the performance of MASTER-seq. It should further be emphasized that even when MASTER-seq is combined with an m6A-IP, the cleavage efficiency metrics that are assessed by MASTER-seq are orthogonal to the coverage metrics that are taken into account in m6A-seq.

We then examined m6A levels at 199 quantifiable ACA sites from the catalog of previously identified putative m6A sites that had been computationally inferred by searching for the nearest methylation consensus motif in the proximity of m6A-seq peaks (‘m6A-seq sites’) (Schwartz et al., 2013). Strikingly, cleavage efficiency scores at m6A-seq sites were strongly reduced in the WT samples compared to the ime4Δ/Δ samples, consistent with the presence of methylation at these sites. Moreover, applying m6A-IP prior to MASTER-Seq resulted in even greater reduction of the cleavage efficiencies in the WT samples but did not impact their counterparts in the ime4Δ/Δ samples (Fig. 2A-B). These results thus demonstrate the ability of MASTER-seq to orthogonally validate putative m6A-sites at single-nucleotide resolution.

**Figure 2.**
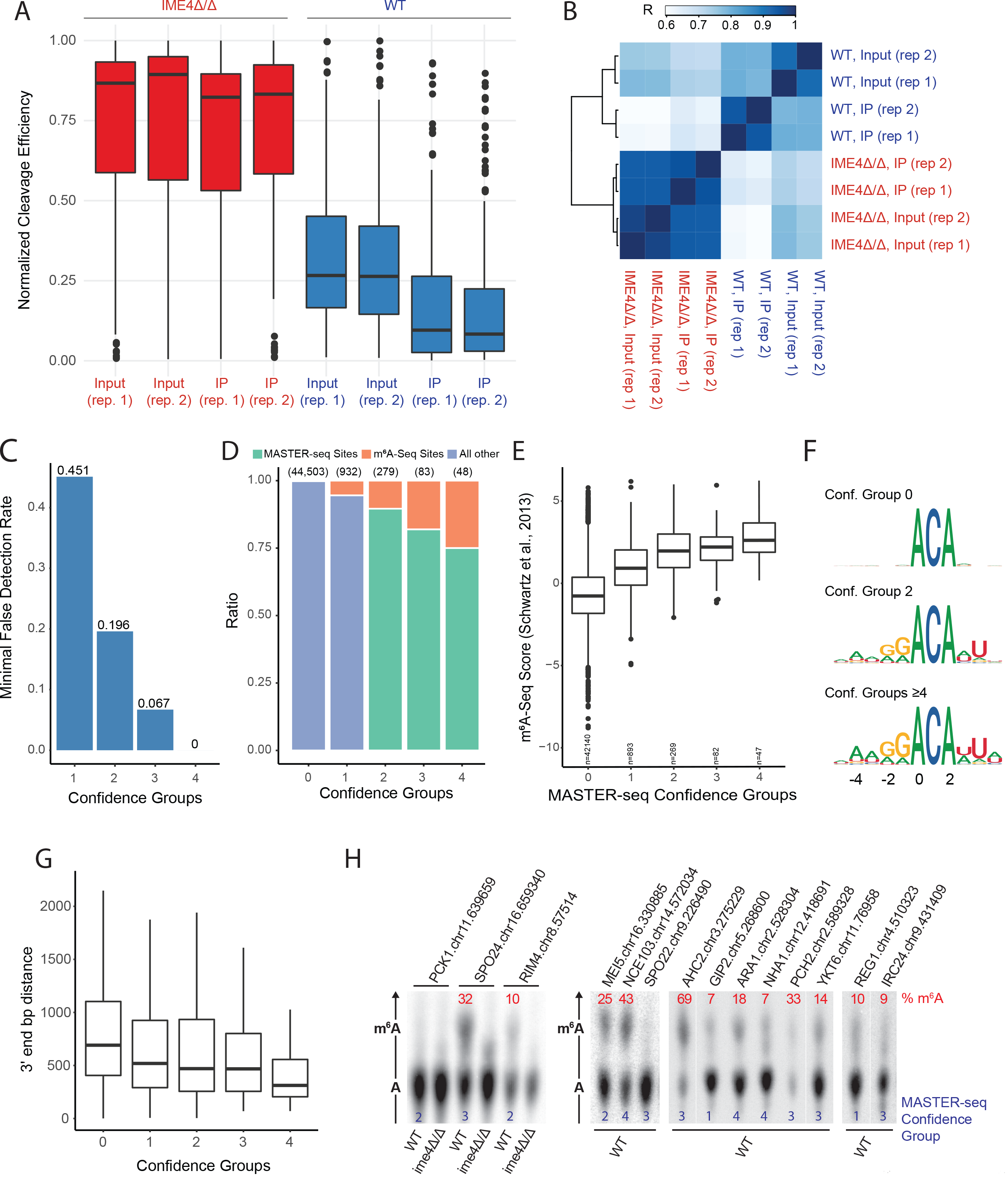
MASTER-seq allows orthogonal validation and discovery of m6A sites. **(A)** Previously reported m6A-seq sites show decreased cleavage efficiencies in WT strains compared to IME4Δ/Δ strains. Normalized cleavage efficiency distributions for “m6A-seq Sites” in replicates with and without m6A-IP selection treatment (Input and IP, respectively). **(B)** Samples correlation. A heat map showing the pairwise correlation between samples, and clustering by similarity to each other. **(C)** Empirical false detection rates per confidence group. **(D)** Distribution of m6A-seq sites across the confidence groups defined via MASTER-seq. **(E)** Distribution of m6A-seq scores from (Schwartz et al. 2013) by confidence groups. **(F)** Sequence logs for sites identified via MASTER-seq, shown separately for sites in confidence group 0 and below, 2 and 4 and above. **(G)** Sites in higher confidence groups are closer to the end of the transcript. Distributions of 3’ end distances by confidence group. **(H)** SCARLET based readouts of methylation levels at each of the indicated sites.

We next sought to detect m6A sites *de novo* using MASTER-seq. We developed an approach relying on three comparisons of cleavage efficiencies: (1) between WT and ime4Δ/Δ Input samples, (2) between WT and ime4Δ/Δ m6A-IP samples, and (3) between m6A-IP and Input (non-m6A-IP) WT samples. For all three comparisons, a true m6A site is expected to have lower cleavage efficiencies in the former condition than in the latter. We assembled a database of 45,845 quantifiable ACA sites across the yeast transcriptome, for which we also had sufficient coverage under the surveyed conditions (‘Methods’). We then classified each of these sites into confidence groups, integrating the number of comparisons in which they scored significantly together with the effect sizes and the P values of these comparisons (Methods, **Fig. S2A**), whereby confidence group 0 is the lowest confidence group and 4 the highest. Remarkably, 410 sites were detected in confidence groups 2 and above, out of which only 56 are part of the previous catalog of m6A-seq sites. The following lines of evidence were used to assess the validity of the *de novo* detected sites (‘MASTER-seq sites’), and to conclude that sites detected in the higher confidence groups (two and above) are particularly highly enriched for bona fide m6A sites: **(1)** We estimated a minimal bound on the empirical false detection rate for each of the confidence groups, on the basis of the number of significant hits in the reverse comparisons (i.e. sites with enhanced cleavage in the WT strain compared to deletion, or in input sample compared to IP) **(Fig. S2A)**. While confidence group 1 (i.e. sites scoring significantly only in one comparison) was associated with a substantial false detection rate, confidence groups 2, 3 and 4+ were associated with minimal false detection rates of ~20%, ~7%, and 0% respectively (Fig. 2C). **(2)** Higher-confidence groups were increasingly enriched in ‘m6A-seq sites’, that had been identified on the basis of peak enrichment (Fig. 2D) **(3)** For each ACA harboring site in the genome an ‘m6A-seq score’ was calculated, on the basis of m6A-seq data from (Schwartz et al., 2013), which quantifies the enrichment in coverage at a site in the IP sample in comparison to the Input sample. We found that sites in increasingly higher confidence groups also showed higher ‘m6a-seq score’ levels, providing strong orthogonal evidence for the validity of the sites (Fig. 2E). Importantly, such enrichment was clearly evident also when performing this analysis only on sites exclusively identified via MASTER-seq and not by m6A-seq (**Fig. S2B**). **(4)** Sites in high confidence groups harbored the same sequence motifs (Fig. 2F) and increasing enrichment towards the transcript 3’ end (Fig. 2G), as was previously reported for m6A-seq sites in yeast (Schwartz et al., 2013). **(5)** m6A-seq and MASTER-seq sites showed similar temporal dynamics across a meiosis time-course (see Fig. 4, below). **(6)** To directly validate the presence of m6A at sites identified by MASTER-seq, we applied SCARLET, a cleavage and ligation-based approach interrogating m6A levels at individual sites directly by radiolabeling and thin layer chromatography (Liu et al., 2013), a low throughput method currently serving as the gold standard in the field. We were able to obtain informative readouts (see **Supplementary Note 1**) for 14 sites that had been exclusively identified via MASTER-seq. These included 2 sites from each of confidence groups 1 and 2, and 5 and 3 sites from confidence groups 3 and 4, respectively. We were able to validate the presence of m6A at levels ranging from 7% to 69% at 12 of these 14 sites (Fig. 2H). At the two remaining sites the observed m6A signal was not appreciably different from background, indicating either no or very low methylation (for further analyses the m6A level at these sites was assumed to be 0%). Collectively, these analyses provide multiple levels of orthogonal support to the newly detected sites in this collection and demonstrate that the false detection rate, in particular in confidence groups 2-4, is low.

MASTER-MINE allows us, for the first time, to use an orthogonal approach to estimate the sensitivity and false detection rate of m6A sites using m6A-IP. The fact that, even in the highest confidence groups, ~4 fold more novel than known sites are detected using MazF based cleavage demonstrates that the antibody-based approach - in their combined experimental and computational implementation - had dramatically underestimated the number of methylation sites. We also estimated the false-detection rate of antibody-based approaches, on the basis of sites that had been detected by m6A-seq but were binned into low confidence groups in MASTER-seq. Of note, sites classified into the low confidence groups comprise both sites that are not methylated (true negatives) in addition to sites that are methylated but for which we lack the statistical power to assign them into higher confidence groups. Nonetheless, we observed that sites that had been called by m6A-seq but had been assigned into low confidence groups tended to be substantially more distant from an m6A consensus site than their counterparts in higher confidence groups. This analysis allowed us to conservatively estimate a minimal false detection rate of 11.3% (see **Supplementary Note 2 for full analysis, Fig. S2C**). MASTER-Seq thus allows us to establish that m6A-IP based approaches are substantially limited in their sensitivity, and also suffer from non-negligible false detection rates.

### M6A deposition and stoichiometry are hard coded into the yeast genome

We next sought to understand the ‘m6A code’, i.e. to explore the extent to which methylation *stoichiometries* can be predicted. Towards this goal, we first sought to identify the optimal *quantitative* measure of m6A levels. If MazF cleavage were 100% efficient at unmethylated sites, all ACA sites in an ime4Δ/Δ strain (which lacks m6A) should be cut at 100% efficiency. However, inspection of the distribution of this metric revealed that cutting by MazF is variable from one site to another (Fig. 2A). One factor appearing to contribute to this is secondary structure of RNA: Cleavage efficiency scores correlated significantly with predicted stability of secondary structure at the region surrounding the ACA site (R = 0.3, Permutation test (10,000 permutations) P < 1×10^−3^), consistent with previous observations that MazF is biased towards cleavage of single-stranded RNA (Zhang et al., 2003) **(Fig. S3A)**. As a quantitative metric of RNA methylation, we thus introduced a ΔCleavage-efficiency metric, capturing the *difference* between MazF cutting in a WT strain, in comparison to an ime4Δ/Δ, under the assumption that this difference will eliminate effects originating from distinct baseline cutting levels at different sites. Indeed, ΔCleavage-efficiency lost most of the correlation with secondary structure (R=0.1, Permutation test (10,000 permutations) P < 1×10^−3^) **(Fig. S3B)**.

The quantitative performance of ΔCleavage-efficiency is supported by several critical lines of evidence. First, and most importantly, ΔCleavage-efficiency levels correlate highly with SCARLET based quantitations, which serve as a gold standard for m6A quantitation (Spearman Rho = 0.79, P = 1×10^−3^, Fig. 3A). Raw cleavage efficiencies also correlated highly, but slightly more poorly, with SCARLET quantitations (Rho = 0.78, P = 1×10^−3^). Second, we observed a significant correlation between ΔCleavage-efficiency and the m6A-seq scores (R = 0.38, Permutation test (10,000 permutations) P < 1×10^−3^), which are derived based on orthogonal measurements **(Fig. S3C)**. Raw cleavage efficiencies correlated slightly more poorly with m6A-seq scores (R = −0.37, Permutation test (10,000 permutations) P < 1×10^−3^, **Fig. S3D**). ΔCleavage-efficiency also correlated better with the fold-change in enrichment in coverage upon IP in the WT strain with respect to the coverage upon IP in the ime4Δ/Δ strain (R=0.44 and R=-0.41, respectively, **Fig. S3E-F**). These data suggest that that while both metrics capture methylation stoichiometry, ΔCleavage-efficiency provides a slightly more reliable relative measurement of methylation levels.

**Figure 3.**
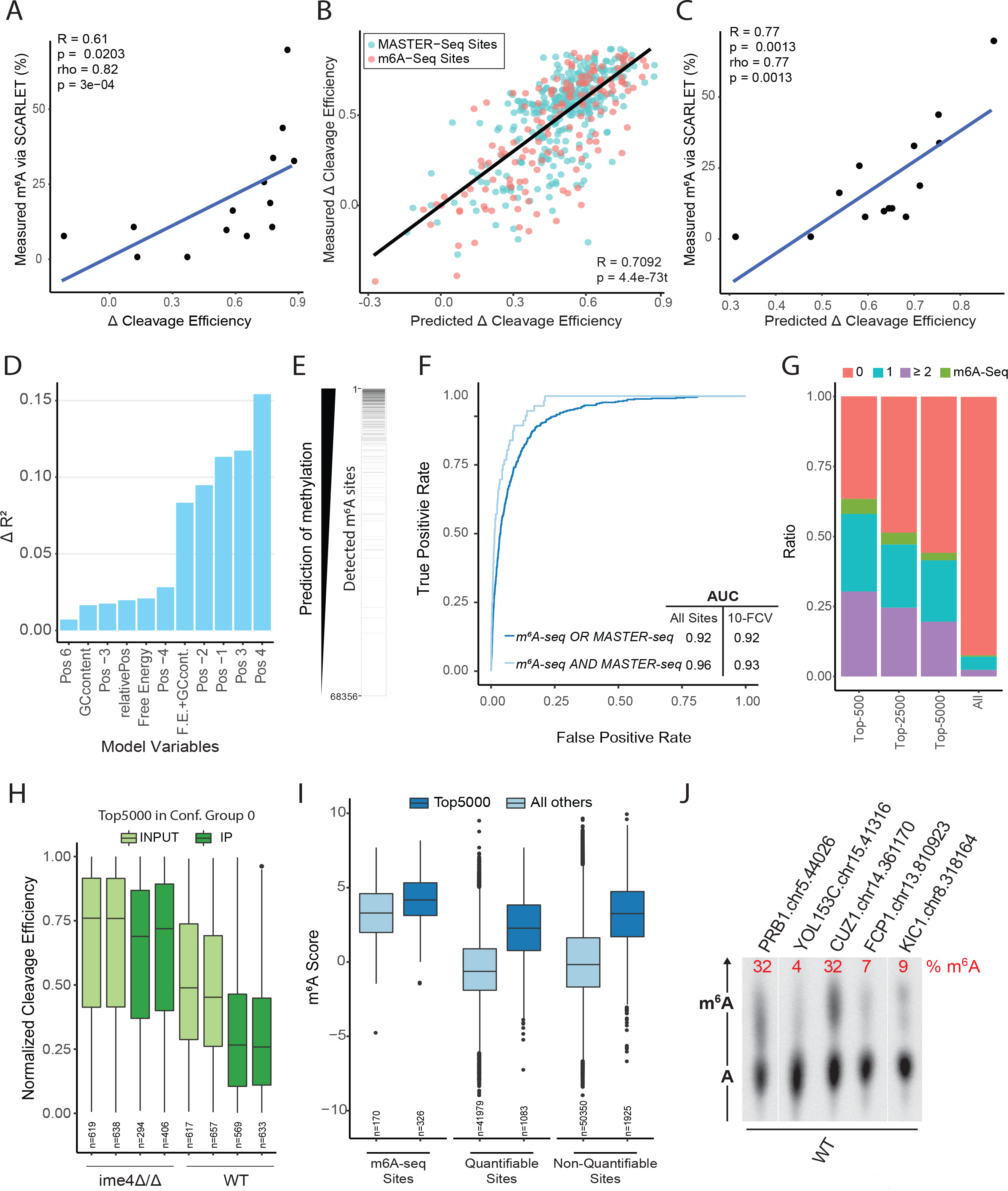
M6A presence and levels are hard-coded ‘in cis’. **(A)** Correlation between the ΔCleavage-efficiency (X axis) and the SCARLET quantitations (Y axis) for 14 sites that were interrogated via SCARLET. **(B)** Variability of methylation levels can be predicted via local sequence and secondary structure information in both new and previously detected sites. Predicted methylation stoichiometry, via a linear model, are plotted against their ΔCleavage-efficiency. A regression line fitting all the values is shown in black. **(C)** Correlation between the linear-model *predicted* cleavage efficiency (X axis) and the SCARLET quantitations (Y axis) for 14 sites that were interrogated via SCARLET. **(D)** Quantification of the relative contribution of each of the indicated variables to the performance of the stoichiometry model. The bar plot depicts the difference in R^2^ from the full model when removing each of the variables in a 1-in-1-out fashion. **(E)** Visual depiction of the predictions made by the model. All RRACA consensus sites are sorted based on their predicted score. A black bar (right) denotes whether methylation was identified at the site (either using m6A-seq or MASTER-seq). **(F)** ROC curves of two predictors of methylation deposition for sequences with the RRACA consensus motif. Both predictors were also tested in a 10-Fold Cross-Validation (FCV) setting and the resulting AUCs are shown. **(G)** For each top-n predictions by the model (whereby n = 500, 2500 and 5000), the distribution of confidence groups among this set of sites is depicted (note that this analysis was performed only for the subset of *quantifiable* sites). In addition, if the site was identified via m6A-seq (but not MASTER-seq) this is indicated in a separate color. For comparison, these numbers are also shown for all sites, serving as a background. **(H)** Quantifiable Top-5000 scoring sites show decreased levels of cleavage in the WT compared with the deletion strain, and further reductions upon m6A-IP. The boxplots indicate the distribution of cleavage efficiency (Y axis) across WT and ime4Δ/Δ yeast cells, either in the presence or absence of m6A-IP. **(I)** Top-5000 sites show extensive enrichment upon m6A-IP in comparison to all remaining sites. Results are shown separately for quantifiable sites, non-quantifiable (via MASTER-seq) sites and which are hence identified exclusively based on the computational model (in Top-5000). As a control, distributions are also shown for sites identified by m6A-seq. Note that the extent of enrichment upon IP is similar across the three sets of sites. **(J)** SCARLET readouts for 5 sites that were non-quantifiable via MASTER-seq, and were predicted purely based on the logistic model.

**Figure 4.**
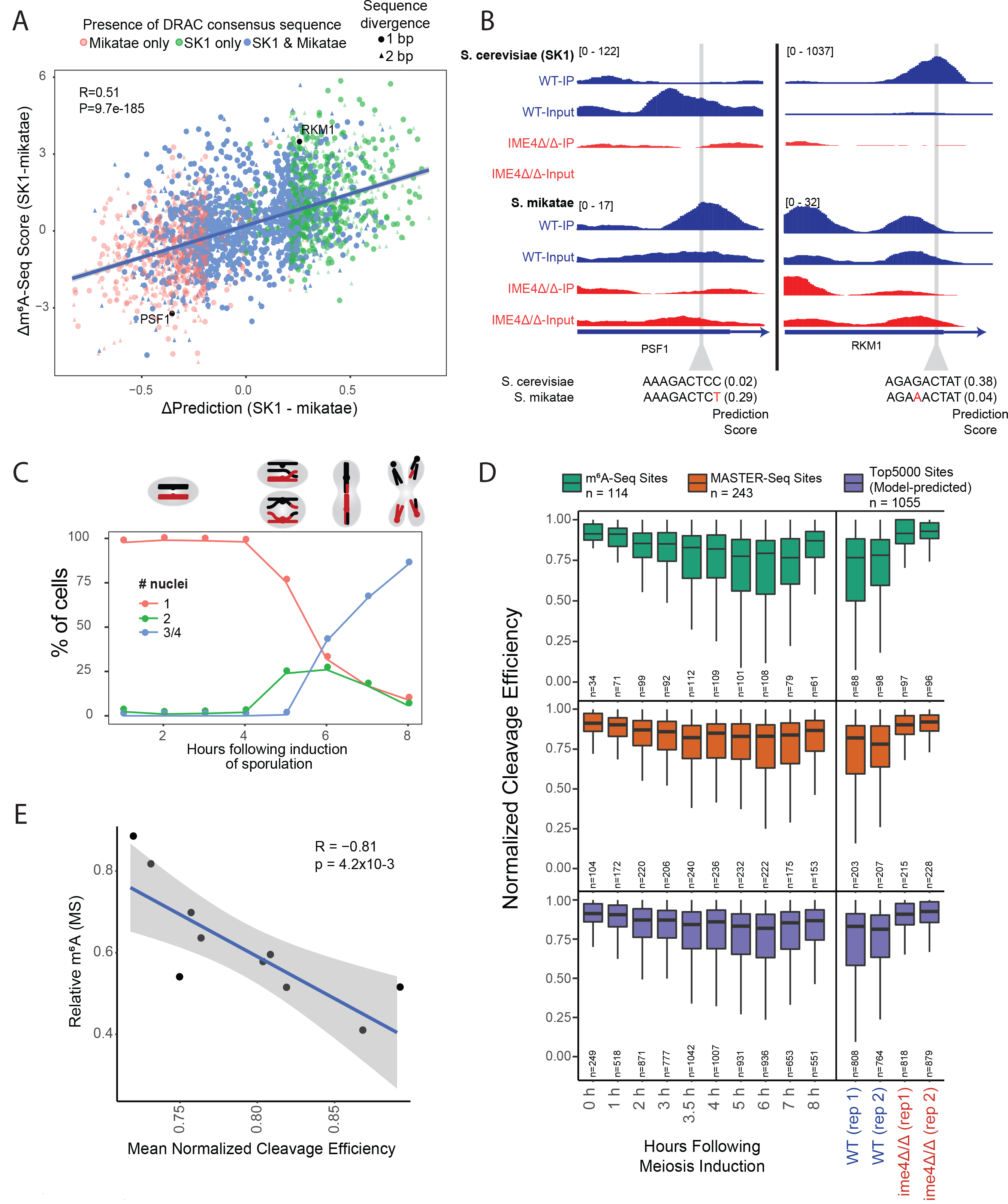
Predictable acquisition and loss of m6A in evolution, and global m6A dynamics in meiosis. **(A)** Scatter-plot depicting the correlation between the difference in the predicted strength of orthologous m6A consensus motifs between SK1 and *S. mikatae* (ΔPrediction, X axis) and the difference in m6A-seq scores at these sites between SK1 and *S. mikatae* (Δm6A-seq score, Y axis). Each point (reflecting an individual site) is plotted as a circle if the sequence divergence in the 9 bp window centered around the methylation site is 1 bp (i.e. the core motifs differ by only a single bp) or triangle is the divergence is 2 bp. The color of each point reflects whether the core sequence motif - defined as presence of a DRAC core - is present only in *S. mikata*e (red), only in *S. cerevisiae* (green) or in both species (blue). The two labelled points indicate the two sites plotted in panel (B). **(B)** Visualization of coverage tracks - obtained from m6A-IP experiments - in both SK1 and *S. mikatae*. Shown are two sites, selected from panel A. The left panel depicts differential methylation in the PSF1 gene between *S. cerevisiae* and *S. mikatae*, linked - in a predictable manner - with changes in the base composition at position +4, leading to absence of methylation in *S. cerevisiae* but presence in *S. mikatae*. The right panel depicts an opposite example, in which methylation is present in *S. cerevisiae* but absent in *S. mikatae*, linked to changes in the composition of position −1. **(C)** Meiotic nuclear divisions kinetics assayed by DAPI DNA staining. Samples from WT SK1 cells (SAy821) were harvested at indicated time points following induction of meiosis, stained with DAPI and the number of nuclei observed for each cell were manually counted. Represented are the percentage of single, double, triple or tetra nuclei cells counted for the different time points. **(D)** Meiotic m6A dynamics measured with MASTER-seq. Distributions of normalized cleavage efficiencies of three different groups of m6A sites: m6A-seq sites, MASTER-seq sites and Model-predicted sites. In parallel, measurements were derived 6 hours following induction of meiosis for ndt80Δ/Δ mutants (SAy841) and for double mutants of ndt80Δ/Δ and IME4Δ/Δ strains (SAy966), as positive and negative controls. **(E)** Correlations between mass-spectrometry-based quantifications of the m^6^A levels and mean normalized cleavage efficiencies, derived from MASTER-MINE.

We next trained a simple linear model using the base identity of 4 bp upstream (positions −1 to −4) and 4 bp downstream (position 3 to 6), of the ACA mazF cutting consensus sequence (note that positions 1 and 2 are fixed), alongside three features capturing: the GC-content, the propensity towards secondary structure (predicted ‘free energy’), and the relative position within the gene. We trained this model based on sites identified exclusively via MASTER-seq, and used the sites identified via m6A-seq as a validation set. The linear model yielded excellent agreement with the MASTER-seq derived ΔCleavage-Efficiencies, not only in the training set but also in the validation set (R^2^ = 0.48 in both cases) **(Fig. S3G)**. A model based on both MASTER-seq and m6A-seq sites achieved an R^2^ = 0.503 (Fig. 3B). As an independent validation of the model, we applied the model to the set of sites measured via SCARLET, and obtained an excellent agreement between the predictions of the model and the measured values (R=0.78, P = 1×10^−3^, Fig. 3C). Critically, training an identical model on the same sites but using m6A-seq scores (instead of ΔCleavage-Efficiencies) explained much less of the variance (R^2^=0.11) when applied to the same training-validation scheme, reflecting the reduced quantitative nature of m6A-seq **(Fig. S3H)**. These findings both establish the predictive power of the model and further support the validity of the novel set of sites. Our results thus strongly suggest that in yeast, m6A stoichiometries are primarily dictated in ‘cis’ via a simple code, embedded in the sequence and structure at the modified site.

To identify and rank the factors contributing to the performance of the model, we examined the coefficients assigned to them by fitting the model **(Table S1)**, and assessed the relative contribution of the variables in the model by removing in a 1-in-1-out fashion the variables, and calculating the difference in the resulting R^2^ from the original R^2^ (ΔR^2^) (Fig. 3D). Interestingly, two of the most relevant variables, in order of ΔR^2^, were the identity of the nucleotides at position +4 and at position +3, with respect to the consensus signal; with a strong positive effect on stoichiometry for a ‘U’ at the former, and a strong negative one at the latter for the presence of a ‘G’. The fact that these two positions are of high predictive power for m6A methylation stoichiometries is of particular interest given that both of them lie outside of the classical, well characterized, DRACH motif, suggesting that, at least in yeast, critical determinants of m6A levels are present outside of the classical motif. These two positions were followed by the identities of position −1 and −2, in both of which ‘G’ is particularly favored, followed by ‘A’, in line with previous observations in yeast and in human (Dominissini et al., 2012; Meyer et al., 2012). Other features, with reduced contribution, included lack of secondary structure at the region surrounding of the modified site along with GC content, also consistent with previous observations, and relative position within the gene, with a bias towards 3’ end (Schwartz et al., 2013). These results thus suggest that nearly 50% of the variability of m6A levels from one site to another is determined primarily via local sequence, with minor contributions from secondary structure and the proximity of the site to the end of the gene.

Given that the stoichiometry of m6A appeared to be to a large extent hard-coded, we next inquired whether m6A presence was similarly hard-coded and hence predictable. For this task, we generated an ultra-high confidence set of m6A sites, comprising all sites identified both in m6A-seq and via MASTER-seq (defined as sites in confidence groups ≥ 2) and harboring an RRACA consensus motif (R=A/G). Remarkably, a logistic classifier trained with these sites as positives and all remaining transcriptomic RRACA sites that had not been flagged as potential m6A sites as negatives, was able to achieve an area under the curve (AUC) of 0.96 (0.93 in a 10-fold Cross Validation setting), indicating an excellent ability to discriminate between methylated and non-methylated sites (Fig. 3E-F). A model generated on the basis of sites defined via m6A-seq *or* MASTER-seq had similar performance (AUC=0.92) (Fig. 3E-F). Of note, the relative weights assigned by the linear and by the logistic classifier were overall very similar **(Table S1, S2)**. Consequently the predictions yielded by the two models were highly correlated (rho = 0.9), and the relative contribution of different variables were similar **(Fig. S3I)**.

We next sought to assess the ability of this model to detect sites *de novo* at a genome-wide level. We applied the above-derived logistic model to each of the 68,356 RRACA sites in the yeast transcriptome **(Table S3)** and examined the 5000 sites with the highest scores (Top-5000). Remarkably, nearly 50% of the quantifiable sites, were classified in confidence groups 1 and upwards, a massive enrichment with respect to the background (7.6%) (Fig. 3G & **Fig. S4A**). Note that the model was trained on sites in confidence groups ≥ 2, hence the enrichment for confidence group 1 (which is enriched 4-fold more than expected **(Fig. S4A)** cannot be due to overfitting. As indicated above, many of the sites in confidence group 1 and the vast majority of sites in confidence group 0 do not reflect truly methylated sites. However, examination of the cleavage efficiencies in the sites forming part of the top-5000 showed that even when these sites were assigned to confidence group 0, they showed strong evidence of decreased cleavage in WT compared to deletion (Fig. 3H), and a substantial enrichment upon m6A-IP, compared to Input (Fig. 3I, **Fig. S4B**). Moreover, also the *non-quantifiable* sites forming part of the top-5000, i.e., the set of sites for which we had been unable to obtain any measurements using MASTER-seq and which had been predicted exclusively based on the model, were also dramatically enriched upon m6A-IP, in fact to a very similar extent as the ‘m6A-seq’ sites that had originally been detected (Schwartz et al., 2013) (Fig. 3H), demonstrating the ability of the model to *denovo* detect m6A sites. Finally, the set of top-5000 sites also showed similar temporal dynamics in a meiosis experiment as the well established m6A-seq sites (see below and **Fig. S4C**). These results thus strongly suggest that a substantial proportion of the top-scoring predicted sites – even the ones corresponding to classes 0 and 1 – are likely truly methylated. Their assignment to lower confidence groups likely reflects the lack of statistical power to classify them into higher bins, most probably due to their decreased levels of expression, which in turn result in lower levels of coverage at the single-base level (**Fig. S2J**).

Finally, to orthogonally validate this genome-wide model, we selected 5 sites from the top-5000 predictions by the model. Importantly, all of these sites were non-quantifiable via MASTER-seq (as they lie within very close proximity to both upstream and downstream ACA sites) and do not form part of the ‘m6A-seq sites’ (Schwartz et al., 2013). As such, the only indication for the presence of m6A at these sites was the fact that they were predicted by the model. Remarkably, we were able to validate the presence of methylation at all 5 sites via SCARLET, at stoichiometries ranging from 4 to 32% (Fig. 3J), demonstrating the power of this model and the predictable pervasiveness of m6A across the yeast transcriptome.

### Single point mutations are sufficient to drive predictable loss and acquisition of methylation sites across evolution

Our findings that m6A is dictated, to such a high extent, in ‘cis’ suggests that changes in sequence - as occur naturally, throughout evolution - should give rise to predictable loss and acquisition of methylation sites. In this sense evolutionary divergence serves as a natural sequence perturbation experiment. To investigate this, we applied the above derived logistic model to all DRAC containing sites in two yeast species: *S. cerevisiae* and *S. mikatae*. We identified all sites predicted to undergo methylation in either of the species, and calculated a Δprediction score, capturing the difference in their predicted likelihood of undergoing methylation in *S. cerevisiae* versus *S. mikatae*. We then made use of available m6A-seq datasets in meiosis for both organisms (Schwartz et al., 2013), and calculated for each site the Δenrichment score, defined as the difference between the m6A-seq score in *S. cerevisiae* and *S. mikatae*. To simplify the interpretation of this comparison, we limited our analyses to cases where the 9 bp methylation consensus window diverged by at most two base pairs between the two species. We observed a striking positive correlation between the Δprediction and Δenrichment scores (R=0.51, P=9.7e-185, Fig. 4A), indicating that loss or acquisition of a methylation site can occur, in a predictable manner, through single point mutations occurring across evolution. Two examples of differential single point mutations occurring between *S. cerevisiae* and *S. mikatae* and leading to predictable differential methylation profiles between the two are depicted in Fig. 4B.

Collectively, these results thus demonstrate that both the presence and the levels of m6A are to a large extent dictated in ‘cis’ via a highly predictable code, and that this code both defines the methylation landscapes within cells, and across evolution.

### Quantitative interrogation of m6A levels in a dynamic response

The inability to quantify m6A levels has been particularly limiting in the context of dynamic cellular and/or disease-related responses, in which potentially subtle changes in m6A level may play important regulatory roles. To evaluate the ability of MASTER-seq to quantify m6A levels across a dynamic response, we applied it to a densely-profiled time-course following induction of meiosis in yeast (Fig. 4C). We observed a gradual increase in methylation levels up to the six-hours time point, which coincides roughly with meiotic prophase, followed by a reduction in the subsequent time points (Fig. 4D). The median methylation levels derived from MASTER-seq were in strong agreement with relative concentration of m6A in the samples measured via mass-spectrometry (R=-0.8) (Fig. 4E). These results demonstrate the ability of MASTER-seq to resolve quantitative differences in m6A levels in a dynamic process. Remarkably, the measurable set of Top-5000 predicted sites displayed highly similar dynamics to those displayed by m6A-seq and MASTER-seq sites. The fact that this set of sites - the vast majority of which had not been detected using any experimental data, but instead had been predicted exclusively based on the model - obtains comparable dynamics lends strong additional support to the validity of the model for *de novo* prediction of methylation levels. Finally, these results further suggest that while the methylation potential of each site may be encoded ‘in cis’ leading to low potential for *local* regulation of m6A levels, *global* regulation of m6A can be achieved likely through titration of the concentrations of the methyltransferase machinery (see Discussion).

One limitation of m6A-seq - and in particular its cross-linking based derivatives - was that it typically had to be tied to a sequencing readout, and in our hands yielded variable results when coupled with qPCR, likely due to the difficulty in designing a proper ‘normalizing’ gene, and the limited quantitative abilities of m6A-IP. This, in conjunction with the high amounts of material required for m6A-IP, has limited the ability to design a cheap and robust quantitative readout of m6A levels, to be used for example in the context of genetic screens. To test whether MazF based digestion could provide such a readout, we designed MazF-qPCR, a qPCR based assay, with which we targeted two methylation sites in distinct genes (**Methods**). Evaluation of the normalized cutting efficiency at both sites recapitulated the expected patterns of methylation, with levels gradually increasing up to prophase, whereupon they decreased (Fig. 5A). These results thus demonstrate that MazF-qPCR is directly amenable for cheap and robust interrogation using low input amounts (our experiments were done using 50 ng of poly(A) mRNA, ~50 fold lower than the requirements for a typical m6A-IP).

**Figure 5.**
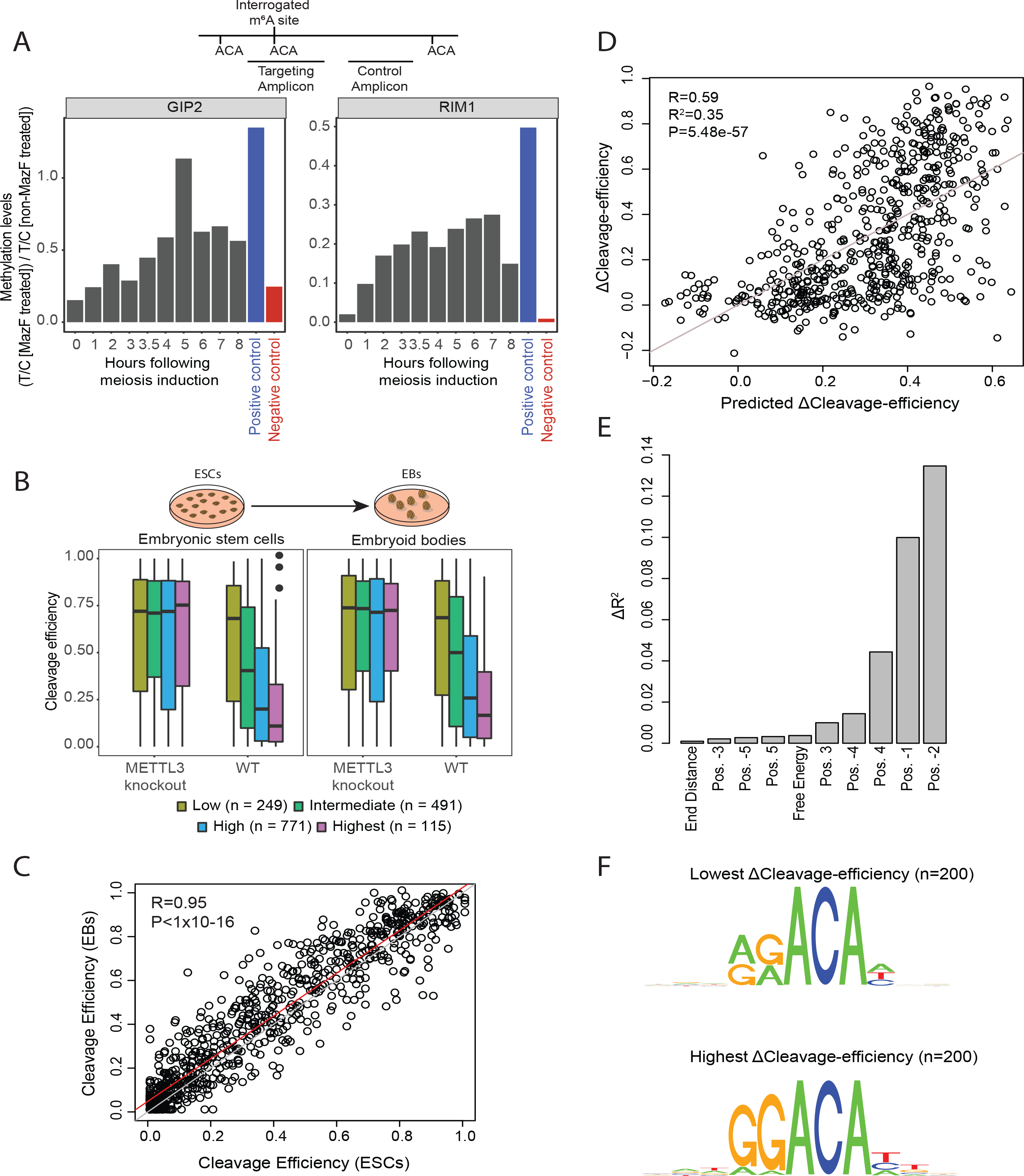
MASTER can be coupled with qPCR readouts and MASTER-seq can quantify methylation within mammalian mRNA. **(A)** MazF-qPCR. Bars represent the methylation levels measured via a qPCR-based assay coupled with mazF digestion for a 10-point meiosis time-course. The levels of a targeted amplicon (‘T’) is measured against a control (‘C’) amplicon in a mazF digested sample, and normalized against a non-digested sample. Positive and negative controls (strains SAy841 and SAy966, 8 hours following induction of meiosis) are also reported. **(B)** MASTER-seq quantification of cleavage efficiencies at m6A sites. Sites are divided into four confidence groups based on the extent to which they were originally predicted to harbor methylation and based on overlap with miCLIP data (Methods). **(C)** Correlation between cleavage efficiencies in embryoid stem cells and in embryoid bodies, shown for all sites of intermediate confidence group and above. **(D)** Correlation between ΔCleavage-efficiencies (Y axis) and the predicted ΔCleavage-efficiencies by the linear model. **(E)** Contribution of different features to the performance of the model, measured by difference in R^2^. **(F)** Sequence motifs, generated on the basis of the 200 sites with the lowest (top) and highest (bottom) ΔCleavage-efficiencies.

### MASTER-seq is applicable to mammalian systems

Finally, we evaluated the applicability of MASTER-seq in mammals. We chose to profile m6A levels in mouse embryonic stem cells (mESCs) and in embryoid bodies (EBs) into which they were differentiated, based on previous findings that knocking out METTL3 in mESCs leads to a differentiation defect that is apparent already within embryoid bodies (Geula et al., 2015). Accordingly, we differentiated either WT or METTL3-KO mESCs into EBs, and applied MASTER-seq to both cell types under both backgrounds, in biological triplicates.

As in yeast, we found that also in this mammalian system MASTER-seq was able to independently validate methylation sites detected via antibody-based methodologies. To evaluate this, we assembled a catalog of quantifiable ACA sites in mouse, divided into low (n=249), intermediate (n=491) and high (n=771) confidence sites, based on the m6A-seq in mouse tissues (Schwartz et al., 2014). In addition, we assigned 115 sites as ‘highest’ confidence if they had also been detected as methylated using miCLIP (Linder et al., 2015). Cleavage efficiencies within both ESCs and EBs progressively decreased in higher confidence groups in WT cells. In contrast, upon METTL3 KO the distributions of cleavage efficiencies across the different confidence groups were roughly identical (Fig. 5B, **Table S4**). We further observed that cleavage efficiencies across the modified sites (excluding low-confidence sites) were highly correlated between ESCs and EBs, demonstrating that no global scaling of m6A levels occurs at the transition from ESCs to EBs (Fig. 5C). These findings demonstrate the applicability of MASTER-seq to mammalian systems, and suggests that the requirement of the methyltransferase complex in the transition from ESCs to EBs probably does not reflect a global redistribution of m6A; Instead, it could potentially reflect a change in how m6A is ‘interpreted’.

Finally, we found that also in mESC, a simple code - primarily capturing the sequence at the modified site - was able to capture 35% of the variability in ΔCleavage-efficiency (Fig. 5D), suggesting that also in mammals a major portion of the m6A signal is hard coded, though possibly to a reduced level than in yeast (see Discussion). Remarkably, we further found that also in mESCs, positions beyond the core DRACH consensus made substantial contributions to the performance of the model. In particular, position +4 - which was the key contributing position to methylation in yeast - also made substantial contributions in mammals (Fig. 5E). In both yeast and mammals, there is a substantial bias for a ‘U’ at this position. Indeed, among the top 200 sites in mESCs ranked by ΔCleavage-efficiency, 50% harbor a ‘U’ at position +4, whereas in the 200 lowest sites no such bias is evident. The principles of the m6A code - and most likely also their mechanistic underpinnings - are thus to a large extent conserved between yeast and mouse.

## Discussion

MASTER-seq adds two critical components which have been lacking for m6A analysis. First, it provides an orthogonal methodology for detecting m6A, and as such allows us for the first time to systematically evaluate the maps derived via antibody-based approaches. These comparisons reveal that the sensitivity of m6A-seq had been severely limited, and that at least 4 fold more sites exist than had been detected using antibody-based approaches. Also the specificity of detection of m6A sites - even after filtering of the 50% of the non-specific peaks that were present in m6A-seq (Schwartz et al., 2013) - remained limited, with a false positive rate of at least 11%. A second critical component introduced by MASTER-seq is an ability to obtain a rough quantification of the stoichiometry of m6A at the detected sites, which has been a key limitation in the field. The inability to quantify m6A levels has precluded unraveling the rules dictating the specificity of m6A deposition, and has imposed major limitations on our ability to explore the extent to which m6A is dynamically regulated both within cells and across diverse stimuli, conditions, and disease states. Two additional advantages of MASTER-seq are its single nucleotide resolution and its applicability to low amounts of starting material. Although in principle variants of m6A-seq exist that separately address either the resolution - using crosslinking approaches - or the latter - by combining m6A-IP with low-input library preparation protocols, no single protocol exists combining the two. We anticipate that MASTER-seq, which we apply here to 100 ng of poly(A) mRNA (~20 fold less than the requirements for a typical m6A-seq experiment), will facilitate the quantitative probing of m6A at single nucleotide resolution in contexts where starting amounts of material are limiting, such as patient-derived samples and lowly abundant tissue/cell types.

A key finding in this study, facilitated by both the enhanced sensitivity and quantitative readout of MASTER-seq, is that the m6A code is remarkably simple. Roughly 50% of the variance in m6A levels in yeast, and 35% in mESCs, is dictated primarily by the sequence immediately flanking the modified site, with relatively minor contribution by the local secondary structure and the relative position within the gene. In mice and yeast, this code includes sequence elements beyond the core DRACH motif, which have likely been missed in previous studies in mammals due to the inability to quantitatively interrogate methylation sites. We estimate that another 15% of the variance in the measurement of m6A levels is technical (in yeast, ΔCleavage-efficiencies correlate with an R^2^ of 0.85). The m6A code elucidated here thus accounts for the majority of non-technical variability in m6A measurements in yeast, and to a substantial proportion of the variability in mammalian systems. Consistently, m6A presence can be predicted *de novo* exclusively based on these features. A corollary of m6A levels being hard-coded is that changes in this code - such as occur during evolution - will lead to changes in m6A levels, either in the form of acquisition or of loss. Indeed, we find that between *S. cerevisiae* and *S. mikatae*, thousands of events involving *predictable* modulation of m6A levels can be observed and inferred directly from the changes in the DNA sequence.

The simplicity of this code has profound implications. Based on work performed in diverse model organism ranging from plants (Arribas-Hernández et al., 2018; Wei et al., 2018), through insects (Haussmann et al., 2016; Lence et al., 2016) to mammalian systems, m6A at different sites is thought to be recognized by diverse readers that lead to differential outcomes, with some readers leading to increased decay, others impacting translation, others impacting export from the nucleus, and additional readers are still awaiting characterization (Dominissini et al., 2012; Edupuganti et al., 2017; Frye et al., 2018; Fu et al., 2014; Gilbert et al., 2016; Meyer and Jaffrey, 2017; Wang et al., 2014; Zhao et al., 2018). In addition, diverse erasers are suggested to be at play, also potentially modulating m6A levels (Jia et al., 2011; Mauer et al., 2016; Zheng et al., 2013). Such complex regulation including multiple factors introduced at distinct stages would be anticipated to result in a complex code, varying substantially from one site to another as a consequence of their propensity to bind and recruit diverse factors. The simplicity of the code suggests that the deposition of m6A is, to a large extent, determined via a single code, across all sites. These findings are consistent with reports that m6A in mammalian systems that the topologies of m6A appear indistinguishable among varying mammalian tissues and responses (Schwartz et al., 2014) and that the m6A content in the nucleus and in the cytoplasm of HeLa cells is essentially identical (Darnell et al., 2018; Ke et al., 2017). While hard-coding of m6A leaves relatively low potential for *local* regulation of m6A levels, it is highly consistent with the *global* regulation of m6A levels, as observed across the meiotic time course (Fig. 4C). Such regulation is most likely achieved through titration of the levels of the methyltransferase complex components, leading to a global scaling of methylation levels. Indeed, mass-spectrometry data across a meiosis time course (Cheng et al., 2018) reveals that Slz1 and Mum2, two critical components of the methyltransferase complex that are required for methylation (Agarwala et al., 2012; Schwartz et al., 2013), are strongly induced at prophase **(Fig. S4D).** This induction likely underlies the massive, global surge in methylation levels at this time point.

Nonetheless, the m6A code that we infer here does not account for 100% of the non-technical variability, and as such additional levels of regulation may be at play. It is possible that the somewhat reduced proportion of the variability explained by our model in mESCs compared to yeast (35% vs. 50%) could be due to the fact that in yeast only a single YTH containing m6A reader is present (Schwartz et al., 2013) and no m6A demethylases have been documented to date. This contrasts with human, which express five different YTH containing readers (Bailey et al., 2017; Dominissini et al., 2012; Ivanova et al., 2017; Shi et al., 2017; Wang et al., 2014, 2015; Zhu et al., 2014), in addition to a growing number of proteins that have been implicated with an ability to read m6A (Edupuganti et al., 2017), and two putative m6A demethylases (Jia et al., 2011; Mauer et al., 2016; Zheng et al., 2013). MASTER-seq provides a fresh, quantitative lens for exploring the sources of variability in methylation levels within cells, and between conditions.

MASTER-seq also suffers from several limitations. *First*, it allows quantification of only a subset of m6A sites that both occur at ACA sites and are within suitable distances of adjacent ACA sites. While this aspect has its disadvantages in terms of not allowing completely unbiased profiling, the reduction in complexity of the RNA fragments - due to cutting exclusively at ACA sites - allows obtaining high-quality signal at lower depth. In this sense, MASTER-seq is analogous to reduced representation bisulfite sequencing (RRBS), which has been instrumental for monitoring and interrogating the roles and dynamics of m5C levels on DNA, by providing a quantitative readout of m5C levels at only a fraction of all methylated sites (Meissner et al., 2005). *Second*, to achieve optimal quantitative abilities, the raw cleavage efficiencies preferably need to be normalized by their counterparts in methylation deficient backgrounds, to allow establishing a ‘baseline’ for each site. Such controls are not readily available for every system. Nonetheless, even in the absence of such methylation deficient controls, *relative* changes in cleavage efficiency can be observed (as was the case in the meiosis time course), and therefore depending on the application, external methylation deficient controls are not strictly required. *Third*, the quantifications obtained via MASTER-seq are tightly connected to the distribution of insert lengths in the sequenced libraries. In our hands, these can differ from one library to another (and from one batch to another) due to minor technical differences in the sample prep, and as a result, we have found that data quality improves upon cross-sample normalization. A *fourth* limitation of MASTER-seq is that MazF is not entirely exclusive to ACA sites, and minor levels of cutting are observed also at ACA resembling sequences, such as ACG or AAA (Fig. 1C), which can result in biased quantifications at some sites. Such biases can potentially be taken into account, and corrected for, in future versions of MASTER-MINE.

MASTER-seq thus provides a highly complementary readout to m6A-seq. Where m6A-seq is unbiased towards a specific motif and of limited quantitative abilities, MASTER-seq is limited to a subset of consensus sequences but offers a quantitative readout. m6A-seq suffers from biases attributable to antibody promiscuity, whereas MASTER-seq is vulnerable to biases originating from the proximity of measured m6A sites to adjacent ACA harboring sites. MASTER-seq provides single nucleotide resolution, that can be further verified via cross-linking based variants of m6A-seq. The availability of MASTER-seq now allows establishing disciplined approaches for benchmarking computational strategies for identifying methylated sites in antibody-based approaches, in a manner seeking to maximize the agreement with this orthogonal method. We anticipate that MASTER-seq will be of high utility for exploring m6A dynamics, functions, mechanisms of action and disease relevance.

## Methods

### Induction of meiosis in yeast

Strain genotypes are detailed in **Table S5**. To induce synchronous meiotic entry, cells were grown for 24 hr in 1% yeast extract, 2% peptone, 4% dextrose at 30°C, diluted in BYTA (1% yeast extract, 2% tryptone, 1% potassium acetate, 50 mM potassium phthalate) to OD600 = 0.2 and grown for another 16 hr at 30°C, 200 rpm. Cells were then washed once with water and re-suspended in SPO (0.3% potassium acetate) at OD600 = 2.0 and incubated at 30°C at 190 rpm. Cells were isolated from SPO at the indicated times and collected by 2 min centrifugation at 3000g. Pellets were snap frozen and stored at −80 for RNA extraction.

### DAPI staining

To observe the progression in cells meiosis, a sample of cells were taken at each time point during sporulation (0 – 8hr) for DAPI staining. Cells were first fixed by Formaldehyde (J.T Baker, UN2209) followed by incubation with DAPI (ThermoScientific, D3571) reagent at 4 °C for 2 hr. Staining was observed by Olympus IX73 Fluorescent microscope system.

### Cell culture

Maintenance of murine WT ESCs or deficient for Mettl3 was conducted as described previously (Geula et al., 2015). Briefly, mESCs expansion was carried out in 500 mL of High-glucose DMEM (ThermoScientific), 15% USDA certified fetal bovine serum (FBS-Biological Industries), 1 mM L-Glutamine (Biological Industries), 1% nonessential amino acids (Biological Industries), 0.1 mM β-mercaptoethanol (Sigma), 1% penicillin-streptomycin (Biological Industries), 1% Sodium-Pyruvate (Biological Industries), 10μg recombinant human LIF (Peprotech). Cells were maintained in 20% O2 conditions on irradiation inactivated mouse embryonic fibroblast (MEF) feeder cells, and were passaged following 0.25% trypsinization. For RNA extraction, cells were grown on Gelatin for three passages in FBS free N2B27-based media (Gafni et al., 2013). Briefly, 500mL of N2B27 media was produced by including: 250 mL DMEM:F12 (ThermoScientific), 250 mL Neurobasal (ThermoScientific), 5 mL N2 supplement (Invitrogen; 17502048 or in-house prepared), 5 mL B27 supplement (Invitrogen; 17504044), 1 mM L-Glutamine (Biological Industries), 1% nonessential amino acids (Biological Industries), 0.1 mM β-mercaptoethanol (Sigma), penicillin-streptomycin (Biological Industries). Naïve conditions for murine ESCs included 10μg recombinant human LIF (Peprotech) and small-molecule inhibitors CHIR99021 (CH, 3 μM-Axon Medchem) and PD0325901 (PD, 1 μM - Axon Medchem) termed 2i. For in vitro embryoid bodies (EBs) formation, roughly 5 million mESCs were disaggregated with trypsin and transferred to non-adherent suspension culture dishes, and cultured in MEF medium (DMEM supplemented with 1% L-Glutamine, 1% Non-essential amino acids, 1% penicillin-streptomycin, 1% Sodium-Pyruvate and 15% FBS – does not contain LIF or 2i) for 9 days. Media replacement was carried out every 2 days.

### mRNA preparation

Yeast total RNA samples were prepared by MasterPure Yeast RNA extraction kit (Lucigen, MPY03100). For mouse ESCs cells, total RNA was extracted using Nucleozol (Macherey-Nagel, 740404.200). Enrichment of polyadenylated RNA from total RNA was performed using Oligo(dT) dynabeads mRNA-DIRECT kit (Thermo Scientific, 61012) for small mRNA amounts. Large mRNA amounts (>1μg) were prepared by GenElute mRNA miniprep kit (Sigma, MRN70). All kits procedures were conducted according to the manufacturer’s protocol.

### MazF digestion

One hundred ng of poly A selected RNA was first heat denatured at 70 °C for 2 min and placed on ice. Each sample was supplemented with 4 µl 5x buffer, 0.8 µl DMSO, 0.5 µl RNase inhibitor (NEB, M0314L) and 20 units of MazF enzyme (TakaRa, 2415A). Reactions were incubated at 37 °C for 30 min and stopped by placing on ice and RNA cleanup.

### RNA barcoding

MazF cleaved RNA was first dephosphorylated with FastAP Thermosensitive Alkaline Phosphatase (Thermo Scientific, EF0654) at 37°C for 30 minutes. Reactions were cleaned of enzymes by adding 3× volume Buffer RLT (Qiagen, 79216) and 1× volume ethanol, precipitating onto SILANE beads, washing twice in 80% ethanol, and eluting in water. We combined dephosphorylated RNA with 20 pmol RNA adaptor 3iLL (**Table S6**), denatured at 70°C for 2 minutes, then snap-cooled by transferring to ice. Ligation reaction proceeded by addition of T4 RNA Ligase 1 (New England Biolabs, M0437M) at 23°C for 75 minutes. Ligated RNA was purified by adding 3× volume Buffer RLT (Qiagen) and 0.85× volume ethanol, precipitating onto SILANE beads, washing twice in 80% ethanol, and eluting in water. Ligated RNA samples were pooled together and the pool was divided into Input sample and IP.

### RNA m6A Immunoprecipitation

m6A-IPs were performed as detailed in (Schwartz et al., 2013). Briefly, RNA was incubated with an anti-m6A antibody (Synaptic Systems, 202003) and protein-G coated beads (Thermo Scientific, 10004D) for 2hr a 4 deg in IPPx1 buffer, followed by magnet pull-down and intensive washing with low salt and high salt buffers. RNA was eluted from antibody using RLT buffer and purified using MyOne Silane Dynabeads (Invitrogen, 37002D) and ethanol. The clean precipitated RNA was then subjected to a second round of IP.

### Library preparation

Libraries were prepared as described in (Safra et al., 2017a, 2017b). Briefly, a DNA primer, complementary to the RNA ligated adapter (**Table S6**), was used for cDNA synthesis by SuperScript lll Reverse transcriptase (Thermo Fisher, 18080093). The reaction was performed according to manufacturer instructions for 1 hr at 50 °C, without heat-inactivation to preserve RNA-cDNA hybrids. The remaining primers left in the reaction were digested by addition of 3μl ExoSAP-IT (Affymetrix, 75001) and incubation at 37 °C for 12 min. ExoSap activity was stopped by addition of EDTA. RNA was degraded by adding 2.5μl of NaOH 1M and incubation at 70 °C for 12 min. The base was neutralized with the addition of 1M HCl, then cDNA was cleaned using SILANE beads as described above. A second adaptor was added to the cDNA by adding 50 pmol 5iLL-22 DNA adaptor (**Table S6**) and ligating with T4 RNA Ligase 1 (NEB, M0437M) at 23°C for 3 hrs. Following clean-up with SILANE beads, we amplified the cDNA library for 13 cycles using barcoded primers (**Table S6**). Amplified libraries were cleaned with AMPure XP beads (Agencourt), quantified using Qubit (Life Technologies) and the distribution of library size was determined using TapeStation (Agilent Technologies).

### LC-MS/MS for determination of m6A/A

To analyze nucleotide composition, 400 ng of double selected polyA RNA fractions were digested with 2 units of P1 nuclease (US biological) at 50 °C in 50 mM ammonium acetate buffer pH 5.3, with 5mM zinc chloride for 2 hours. Nucleotides were dephosphorylated by addition of 5 units of CIP (New England Biolabs) for another 2 hours at 37°C, and then diluted 1:10 in acetonitrile. The samples were dried by acetonitrile evaporation in speedvac. The residue of each sample was re-dissolved in 198μL of 0.01% formic acid. Two μL of 1μg/mL 7-deaza-A was added as internal standard. The mixtures were intensively vortexed (0.5min), centrifuged (21,000rpm; 5min), and passed through 0.22-μm PVDA filters (Millex GV) to 250-μL inserts of LC-MS vials. The LC-MS/MS instrument consisted of an Acquity I-class UPLC system (Waters) and Xevo TQ-S triple quadrupole mass spectrometer (Waters) equipped with an electrospray ion source and operated in positive ion mode was used for analysis of nucleosides. MassLynx and TargetLynx software (version 4.1, Waters) were applied for the acquisition and analysis of data. Chromatographic separation was done on a 100mm × 2.1mm internal diameter, 1.8-μm UPLC HSS T3 column equipped with 50mm × 2.1mm internal diameter, 1.8-μm UPLC HSS T3 pre-column (both Waters Acquity) with mobile phases A (0.01% formic acid) and B (50% aqueous acetonitrile with 0.01% formic acid) at a flow rate of 0.2mL/min and column temperature 25°C. A gradient was used as follows: the column was held at 0%B for 1 min, then a non-linear increase (curve 8) to 35%B from 1 to 18min, then a non-linear increase (curve 8) to 100%B 18-18.2 min, held at 100%B 18.2-19min, back to 0% B 19-20 min and equilibration at 0% B for additional 5min. Samples kept at 7°C were automatically injected in a volume of 1 or 3μL, to get non-saturated A and m6A signals, respectively. Retention times were 9.7, 11.4, 11.8, and 13.8 min for 7-deaza-A, G, A, and N6-Me-A respectively. For mass spectrometry, argon was used as the collision gas with a flow of 0.10mL/min. The capillary voltage was set to 2.67kV, source temperature 150°C, desolvation temperature 400°C, cone gas flow 150L/hr, desolvation gas flow 800L/hr.

Nucleoside concentration was calculated using a standard curve of the relevant nucleotide concentration in each sample. Standard curves included increasing concentration of all measured nucleosides ranging from 0-1000ng/mL that were positioned at the beginning and at the end of each run. All the calculated values for the different nucleosides in each sample fell within the standard curve range. The compounds were detected in positive mode as multiple-reaction monitoring, with the following parameters: 267.1>118.1 and 267.1>135.0 m/z (collision energy CE 57 and 16 eV respectively) for 7-deaza-A, 284.2>152.1 m/z (CE 14eV) for G, 268.1>136.1 m/z (CE 15 eV) for A, and 282.1>123.1 and 282.1>150.1 m/z (CE 40 and 25 eV respectively) for N6-Me-A.

### SCARLET

Site-specific cleavage and radioactive-labeling followed by ligation-assisted extraction and thin-layer chromatography (SCARLET) was carried out as detailed previously (Liu et al., 2013). For each site, 1 µg of mRNA was analyzed using the oligonucleotide pairs listed in **Table S6** for cleavage and ligation together with the previously described 116-mer DNA oligonucleotide (Liu et al., 2013).

### In vitro spike-ins preparation

Two synthetic RNA fragments (IVT1 and IVT2, **Table S6**), each comprising a 103 nt long sequence with a single ACA in the center were in vitro transcribed from dsDNA templates, either in the presence of ATP or N6-methyl-ATP, using MaxiScriptT7 kit (Invitrogen, AM1320). Purified m6A-containing products were serially diluted in non-m6A containing products.

### MazF-qPCR

For the qPCR based readouts of methylation, we designed two primer pairs to interrogate each of two sites. In each case, one ‘test’ primer pair was designed to flank a putative methylation site, and hence upon MazF cleavage will only lead to a product if the site is indeed methylated. The second ‘control’ primer pair was designed to flank an adjacent region in the same gene that did not harbor an ACA site. Readouts were obtained for both pairs either upon MazF digestion or in its absence. The ratio of the abundance of the test versus control primer, in the presence of MazF, was normalized by the corresponding ratio in the absence of MazF treatment. Primer sequences are provided in **Table S6**.

### Read Alignment

Reads were aligned using STAR (V. 2.5.3a) (Dobin et al., 2013), for the reference genome generation step we used the SK1reference genome used by (Schwartz et al., 2013), along with the IVT sequences (Tables S3) as additional chromosomes. Additional parameters used in the alignment step were ‘-- alignIntronMax 300 --alignMatesGapMax 1000’, all other parameters were set by default. Normalization of paired libraries insert size was performed using in-house python scripts, between pairs of INPUT-IP samples. Alignment sorting and indexing were performed separately with samtools (V. 1.3.1) (Li et al., 2009). Single-base coverage was retrieved using in-house python scripts and bedtools (V. 2.26.0) (Quinlan and Hall, 2010), calculating coverage from 3’ and 5’ positions, and fragment coverage (i.e. from the start of read-1 to the end of read-2).

### Cleavage efficiency

Strand specific 3’ positions, 5’ positions, and fragment-coverages were retrieved for all ACA positions using bedtools’ genomecov function and in-house processing scripts. Preliminary cleavage efficiencies were calculated as a ratio of the 3’ read-ends or 5’ read-ends coverage divided by their respective fragment coverage. To identify the minimal distance between an interrogated ACA site and an adjacent ACA site yielding quantitative results, we employed the following scheme for each library: (1) Iterate over all distances between adjacent ACA site from 0 to 200 bp, and compute the pairwise correlation between 5’ and 3’ cleavage efficiencies for all the sites that are within greater distances of adjacent upstream and downstream ACA sites. (2) Using the vector of correlation coefficients in step 1, in a step-forward procedure the ‘optimal ACA distance’ is selected as the one at which the correlation does not increase for the next two increments in ACA distance. Using the resulting distance, a separate calculation is done for downstream and upstream direction separately as defined above, but locking the complementary direction to the optimal distance calculated (**Fig. S1B**). All measurements originating from 5’ or 3’ cleavage efficiencies that were within greater proximity to an adjacent ACA site than the optimal distance were subsequently defined as non-available (N/A) measurements. The final cleavage efficiency was defined as the aggregated cleavage efficiencies originating from all quantifiable 5’ and 3’ cleavage efficiency scores. For the cases of first and last ACA sites in a gene, the missing closest ACA site distance was replaced with the gene start and end coordinates respectively. Of note, as the SK1 transcriptome annotations do not define UTRs we extended each CDS gene coordinates by 150 bp in the 5’ direction and 250 bp in the 3’ direction.

### IP/INPUT differential analysis

IP-m6a enrichment calculations were performed using the ‘edgeR’ package (Robinson et al., 2010). The median of a 50-nt window centered in each ACA sites was used to represent the coverage level of the site, using the previously calculated fragment coverage. Fold-changes were calculated between IP and INPUT treatments in WT strain (m6A-Score), and between WT and IME4Δ/Δ strains in IP treatment.

### Detection of Putative m6A Sites and confidence group assignment

Putative m6A sites were identified using a three-way comparison approach, the comparisons being: (1) sites with less cleavage efficiency in WT than IME4Δ/Δ for INPUT samples, (2) sites with less cleavage efficiency in WT than IME4Δ/Δ in m6A-IP samples, and (3) sites with less cleavage efficiency in m6A-IP than INPUT in WT samples. Due to initial differences between libraries in cleavage efficiency distributions (likely originating from differences in the insert sizes of the libraries, see **Discussion**), a cross-sample quantile normalization was performed using the ‘preprocessCore’ package (Bolstad, 2013). Mean difference, log2 Fold-Change (log2FCH) and t-test p values were calculated for each site in each of the three comparisons, when fragment coverage at the interrogated position was greater or equal to 15 in all the samples in each comparison.

For confidence group assignment, each site was scored: −2, −1, 0, +1, or +2; depending on whether it was associated with a low or high combination of thresholds for effect size and P value. Specifically, for comparisons 1 and 2, the thresholds for associating a site with a +2 score were mean difference of 0.7 and P < 0.01; For a +1 score, the corresponding numbers were 0.5 and 0.05. For comparison 3 thresholds, a +2 score required a log2FCH of 4 and P < 0.01, and a +1 required a log2FCH of 2 and P < 0.05. Scoring significantly for the symmetric but opposite direction would grant a −1 or −2 score respectively **(Fig S2A)**. The scores obtained from the three comparisons were summed for each site resulting in the confidence group assigned to the site. As the number of sites that were assigned to confidence groups 5 and 6 was very small, they were collapsed into confidence group 4. All confidence groups 0 and below were similarly collapsed into the ‘0’ confidence group.

### Methylation models

The variables used for m6A modeling were: (1) binarized nucleotide identity for positions −4 to +6 with respect to the interrogated nucleotide, omitting positions 0 to 2 due to the ACA consensus sequence; (2) local minimum free energy within a 60-nt window centered on each ACA site, estimated via the RNAfold function using parameters `--noconv --noPS -T 30` in the ViennaRNA package (Lorenz et al., 2011); (3) relative position within the gene (in a 0 to 1 scale); and (4) GC content expressed as the percentage of G’s and C’s within the same 60-nt window as minimum free energy.

For the stoichiometry model, we fitted a linear model to the ΔCleavage-efficiency measurements of all the MASTER-seq sites and m6A-seq sites with feature-selected variables. For selection of variables, we performed an exhaustive backward and forward feature selection using the ‘leaps’ package (Lumley and Miller, 2009), and performed a 5-fold cross-validation using in-house scripts to control for over-fitting.

For the binary (logistic) classifier, we modeled methylation fitting a logistic model assigning a positive value for all MASTER-seq sites and m6A-seq sites, and a negative value for all the sites that were in confidence group 0 neither are m6A-seq sites. All variables aforementioned were used without performing feature selection and we performed a 10-fold cross-validation using in-house scripts to control for over-fitting.

### Evolutionary analysis

Pairwise alignments between the SK1 and the *S. mikatae* transcriptomes were performed as in (Schwartz et al., 2013). A custom script was used to identify all DRAC motifs occurring in either of the two transcriptomes, and extracting the sequence environment surrounding it. The binary classifier (above) was applied to each of the identified sites, and for each site the difference between the predicted scores in the two organism was calculated (Δprediction). Sites that lacked a DRAC core (in one of the two organisms) were manually provided a predicted level of 0. Note that the logistic model was only trained on RRACA sites, yet it was applied to all DRAC sites, including both Ts at position −2 and nucleotides other than A at position +2. To expand the model, we treated Ts at position −2 as A (i.e. they received the same weight as an A at this position), and all positions at position +2 were treated as As (we found that limiting ourselves only to RRACA sites did not alter our conclusions, and yet reduced the number of surveyed data points). For the calculation of the respective m6A levels in each of the two organisms, we utilized the available m6A-seq data that had been generated for both of them (Schwartz et al., 2013). For each site in each organism, we calculated the log2FCH in edgeR normalized coverage between IP and Input samples. The Δenrichment score was defined as the difference between this value in SK1 and the corresponding value in *S. mikatae*. For the analysis presented in Fig. 4A-B, we limited ourselves to sites that had at most 2 mutations in the sequence window of 9 bp centered around the methylated site, and that had predicted levels of methylation greater than >0.2 either in SK1 or in *S. mikatae*. In addition, we classified each site based on whether or not a ‘core’ consensus sequence - defined as presence of a DRAC motif - was present in each of the two organisms; This was used only for color-coding each point in Fig. 4A.

### Quantification of cleavage efficiencies in mESCs and EBs

A catalog of quantifiable ACA sites in mouse was obtained from Supplementary Table S2 in (Schwartz et al., 2014). We intersected this dataset with a miCLIP dataset (Linder et al., 2015) obtained for mouse brain (A. Grozhik & S. Jaffrey, unpublished dataset) and defined an ultra high confidence set of sites consistently identified in both. This initial dataset was filtered to retain only (1) sites harboring an ACA, (2) sites within 60 bp from an upstream or downstream ACA sites (and hence quantifiable via MASTER-seq), and (3) sites covered by at least 20 reads (aggregated across all replicates) in at least one measurement.

### Statistical Analysis and plotting

All statistical analyses and visualizations were performed using the R language and the ‘base’ package (Team, 2017), along with supplementary packages: ‘AUC’ (Ballings and Van den Poel, 2013), ‘dplyr’(Wickham et al., 2015), ‘magrittr’ (Bache and Wickham, 2014), ‘tidyr’ (Wickham, 2016), and ‘gtools’ (Warnes et al., 2014). Sequence logos were prepared using the ‘ggseqlogo’ package (Wagih, 2017), most plots were generated using the ‘ggplot2’ package (Wickham, 2010) along supplementary plotting packages: ‘RColorBrewer’ (Neuwirth, 2014), ‘LSD’ (Schwalb et al., 2015), and ‘gridExtra’ (Auguie, 2016).

