## Supplementary Materials for "Deciphering the ‘m^6^A code’ via quantitative profiling of m^6^A at single-nucleotide resolution"

### Deciphering the 'm6A code' via antibody-independent global, semi-quantitative profiling of m6A at single-nucleotide resolution

Miguel Angel García-Campos[1,*], Sarit Edelheit[1,*], Ursula Toth[2], Ran Shachar[1], Ronit Nir[1], Lior Lasman[1], Alexander Brandis[3], Yaqub Hanna[1], Walter Rossmanith[2], Schraga Schwartz[1,#]

[1]Department of Molecular Genetics, Weizmann Institute of Science, Rehovot, Israel 7610001

[2]Medical University of Vienna, Center for Anatomy & Cell Biology, Währinger Straße 13, 1090 Vienna, Austria

[3]Life Sciences Core Facilities, Weizmann Institute of Science, Rehovot, Israel 7610001

* These authors have contributed equally to this manuscript.

**Supplementary Note 1**

In addition to the total of 19 sites at which we quantified m6A levels using SCARLET across the different parts of the manuscript (presented in **Figure 2H** and **Figure 3I**), there were 4 additional sites that we had attempted to quantify, but for which we failed to obtain informative measurements. In all of these cases, TLC yielded very strong signals - inconsistent with the expression level of the genes harboring the putative interrogated m6A sites - that “smeared” substantially across the ‘A’ and ‘m6A’ areas. We interpret these results as likely reflecting some form of non-specific labeling, and these data points were hence not taken into account in the analyses. For completeness, these sites and the oligonucleotides used for their analysis are provided in **Table S6**.

**Supplementary Note 2**

We also sought to estimate the false-detection rate of antibody-based approaches. Of the 199 previously identified quantifiable m6A-Seq sites at ‘ACA’ consensus sequences, 92 were ranked in confidence groups 0. These sites should reflect a mixture of erroneously called sites using the m6A-Seq approach, along with MASTER-seq false negatives, i.e. true m6A sites for which we lacked statistical power to assign them into higher confidence groups. To obtain a *minimal* bound on the erroneously called sites in m6A-Seq sites, we re-analyzed the distribution of the distances between the center of the peaks in m6A-seq data and the called consensus sites across the different confidence groups. Strikingly, this analysis revealed that this distance was almost invariably low (median: 0 nt, maximum: 41 nt) in the high confidence groups, but substantially higher in confidence groups 0 and below (median: 8 nt, maximum: 311 nt). Thus, in the lowest confidence groups, 24 out of 199 (12%) sites were more than 41 nt away from the nearest consensus site **(Fig. S2C)**. Under the conservative assumption that all sites in confidence group 0 with a distance greater than 41 nt (the maximal distance in the highest confidence groups) are false positives, this allows us to estimate a *minimal* false detection rate for all other m6A-seq sites at 11.3% (148 out of 1160).

**Supplementary Figures**

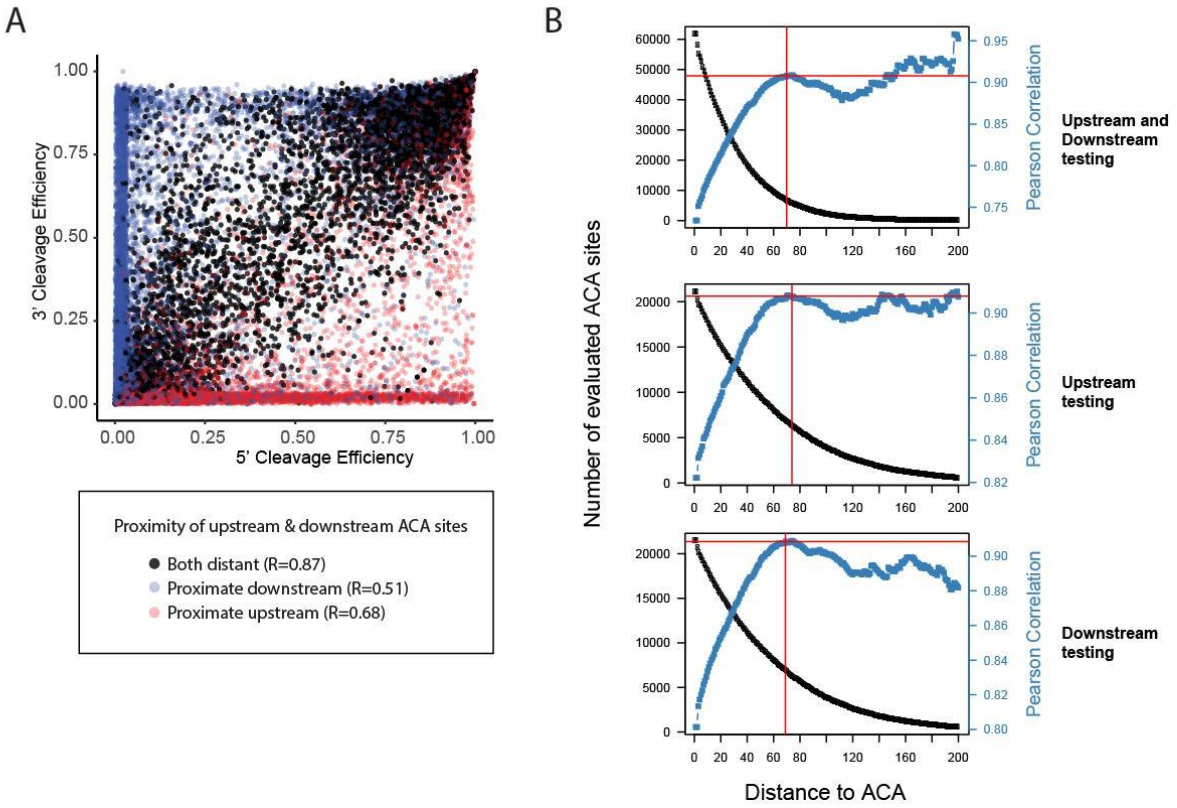


**Figure S1.** Quantitation of m6A signal is dependent on distance to adjacent ACA sites. **(A)** Scatterplot of 3' cleavage efficiency vs 5' cleavage efficiency. Points are color coded based on their proximity with other ACA sites: blue, if the plotted site is too close (≤ 60 nt) to a downstream ACA site; red if it is too close (≤ 60 nt) to an upstream ACA site; and black if it is >60 nt from both closest upstream and downstream ACA sites. **(B)** Minimal ACA distance calculation. Correlations between all 3' and 5' cleavage efficiencies at different minimal ACA distances are shown in blue. The number of evaluated sites at each filtering step is plotted in black. Two red lines mark the point at which correlation stops increasing as minimal ACA distance increases.

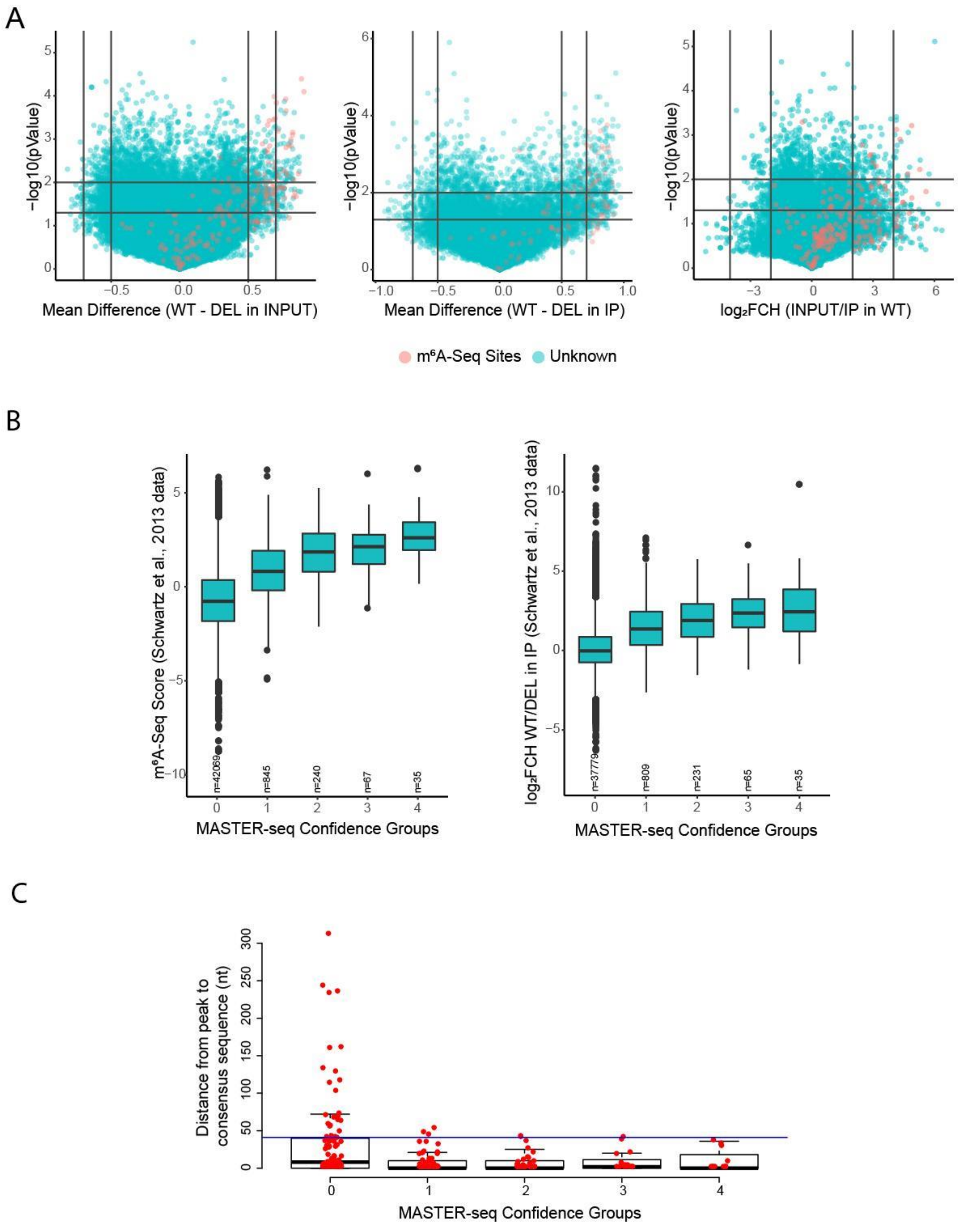


**Figure S2.** *De novo* detection of m6A using MASTER-seq. **(A)** Volcano plots of the three comparisons used in m6A sites *de* novo detection. Vertical and horizontal lines depict the scoring thresholds. X-axis from left to right: 1) Mean difference between cleavage efficiency of each site in WT and IME4Δ/Δ strains in INPUT treatment, 2) Mean difference between WT and IME4Δ/Δ strains in m6A-IP treatment, 3) log2-Fold-Change (Log2FCH) in Input over IP treatments in WT strain. Comparisons 1 and 2 thresholds are (left to right): -0.7, -0.5, 0.5 and 0.7; comparison 3 thresholds are: -4, -2, 2, and 4, respectively. Y-axis: -$\log_{10}$ t-tests p-values; horizontal thresholds are (bottom-up): 0.05 and 0.01 for all plots. **(B)** Boxplots depicting the enrichment of m6A-seq derived scores (Schwartz et al., 2013) in MASTER-seq derived confidence groups exclusively in MASTER-seq *de novo* detected sites. **(C)** Boxplots depicting m6A-seq sites distance from peak to consensus sequence identified in (Schwartz et al., 2013) divided by MASTER-seq confidence groups. A horizontal blue line marks the 41 nt distance threshold, and highlights that confidence group 0, in particular, is associated with a large number of sites that are not within vicinity of a near consensus site.

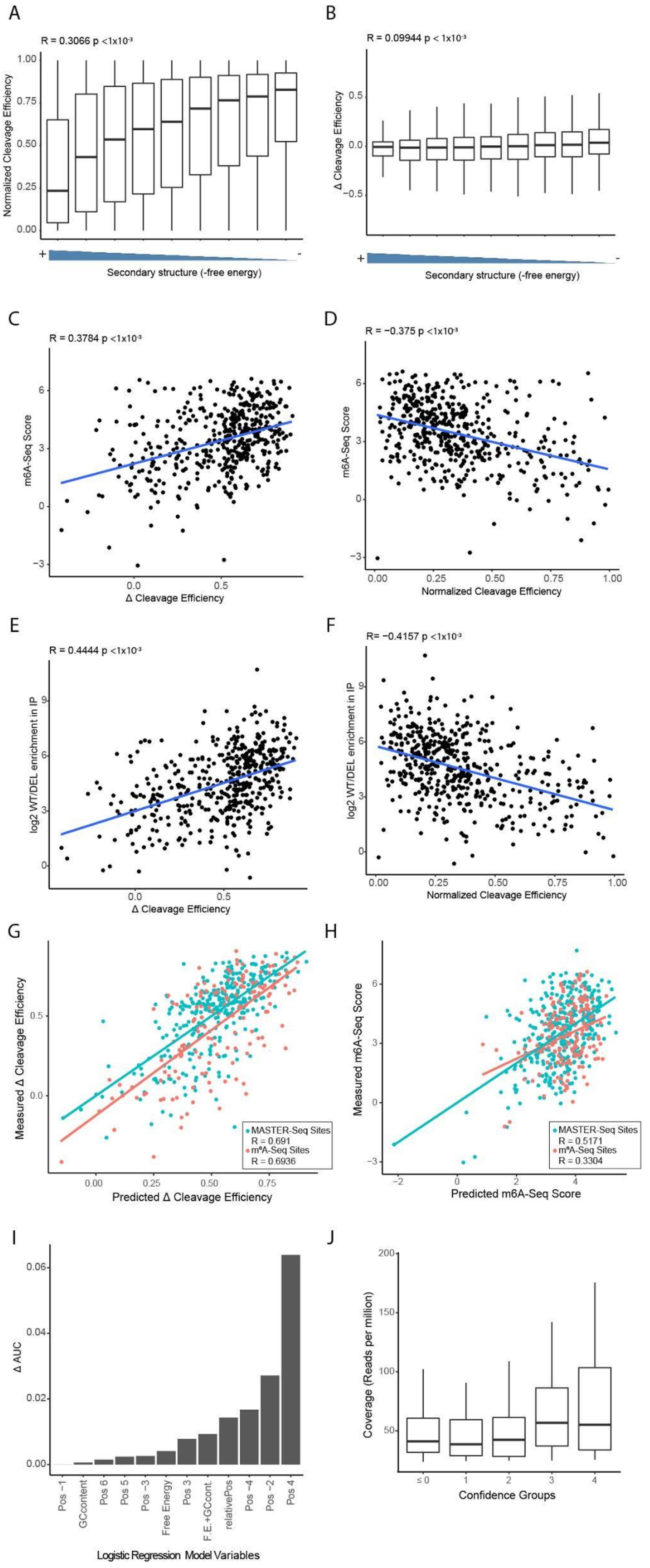


**Figure S3.** M6A is hard-coded. **(A)** Correlation between secondary structure (estimated free energy) and normalized cleavage efficiency measured in IME4Δ/Δ strain. **(B)** Correlation between secondary structure (estimated free energy) and ΔCleavage efficiency. **(C)** Correlation between ΔCleavage efficiency and m6A-seq score. **(D)** Correlation between Normalized cleavage efficiency and m6A-seq score. **(E)** Correlation between ΔCleavage efficiency and $\log_2$ WT/DEL enrichment in m6A-IP. **(F)** Correlation between Normalized cleavage efficiency and $\log_2$ WT/DEL enrichment in IP. **(G)** Correlation between predicted and measured ΔCleavage efficiency. Dots and regression lines are color-coded separately for MASTER-seq and m6A-seq sites. **(H)** Correlation between predicted and measured m6A-seq score, color-coded as in (G). **(I)** Quantification of the relative contribution of each of the indicated variables to the performance of the logistic model. The bar plot depicts the difference in AUC from the full model when removing each of the variables in a 1-in-1-out fashion. **(J)** Coverage in reads per million by MASTER-seq confidence groups.

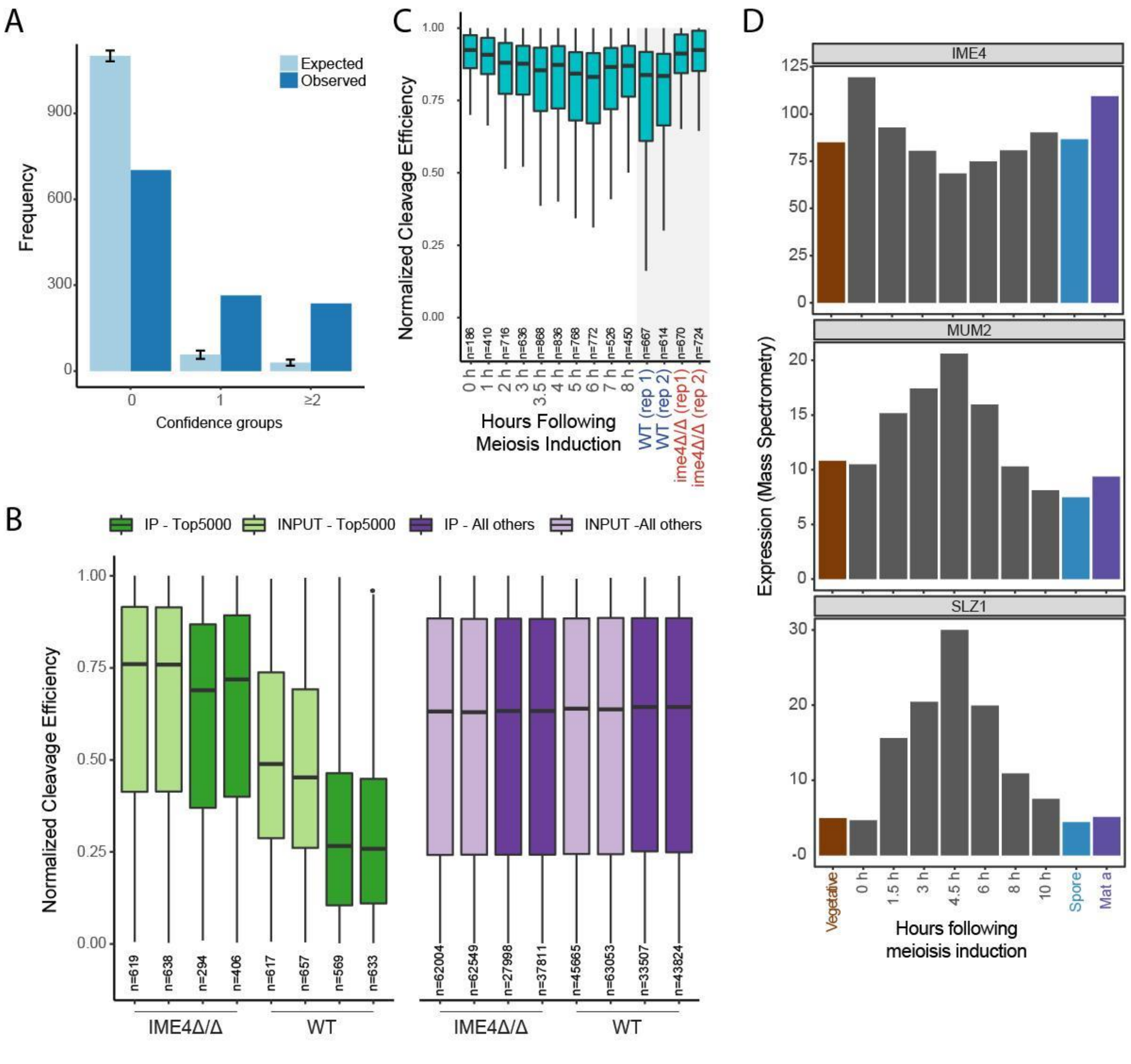


**Figure S4. (A)** Frequencies of expected and observed sites from different confidence groups in the Top-5000 sites set. Expected sites were calculated based on the frequency of each confidence in the entire set of sites (background). Interval bars represent 95% confidence intervals. **(B)** Normalized Cleavage efficiency distributions per sample. Top-5000 MASTER-seq quantifiable sites are shown in green, and background levels of normalized cleavage efficiency are shown in purple. **(C)** Meiotic m6A dynamics in predicted sites measured with MASTER-seq. Distributions of normalized cleavage efficiencies of Top-5000 predicted sites that do not overlap with any of the detected sites (either via MASTER-seq or M$^6$A-seq Sites). **(D)** Mass spectrometry derived measurements of Ime4, Slz1 and Mum2 expression levels across a meiosis time course. Data obtained from (Cheng et al. 2018).

#### Table S1: Linear model coefficients

| | Estimate | Std. Error | t value | Pr(>\|t\|) |
|---|---|---|---|---|
| *Free Energy* | 0.012 | 0.003 | 4.356 | 0 |
| *GCcontent* | −0.612 | 0.158 | −3.875 | 0 |
| *Relative Position* | 0.165 | 0.039 | 4.229 | 0 |
| *POS_−2_G* | 0.262 | 0.035 | 7.388 | 0 |
| *POS_−1_G* | 0.231 | 0.024 | 9.781 | 0 |
| *POS_3_C* | −0.132 | 0.03 | −4.366 | 0 |
| *POS_3_G* | −0.352 | 0.036 | −9.743 | 0 |
| *POS_4_T* | 0.251 | 0.021 | 11.826 | 0 |
| *POS_−4_A* | 0.071 | 0.021 | 3.378 | 0.001 |
| *POS_−3_C* | 0.099 | 0.031 | 3.233 | 0.001 |
| *POS_−2_A* | 0.104 | 0.036 | 2.899 | 0.004 |
| *(Intercept)* | 0.149 | 0.074 | 2.027 | 0.043 |
| *POS_−4_G* | −0.05 | 0.029 | −1.708 | 0.088 |
| *POS_6_C* | −0.047 | 0.027 | −1.707 | 0.088 |
| *POS_3_A* | −0.028 | 0.02 | −1.413 | 0.158 |
| *POS_−3_G* | −0.044 | 0.032 | −1.399 | 0.162 |
| *POS_6_T* | 0.027 | 0.019 | 1.372 | 0.171 |
| *POS_−1_C* | −0.262 | 0.191 | −1.367 | 0.172 |
| *POS_−3_A* | 0.024 | 0.02 | 1.166 | 0.244 |

**Table S1.**
Summary of coefficient estimates for variables in the linear regression model.

#### Table S2: Logistic model coefficients

| | Estimate | Std. Error | z value | Pr(>\|z\|) |
|---|---|---|---|---|
| *POS_-1_A* | −2.5613 | 0.1319 | −19.4224 | 0 |
| *POS_4_A* | −2.0006 | 0.1487 | −13.4558 | 0 |
| *POS_4_G* | −2.0493 | 0.1642 | −12.4784 | 0 |
| *POS_-2_A* | −1.4071 | 0.114 | −12.3419 | 0 |
| *POS_4_C* | −2.4373 | 0.2155 | −11.308 | 0 |
| *Relative Postion* | 2.1542 | 0.2413 | 8.9288 | 0 |
| *POS_3_G* | −1.5221 | 0.2038 | −7.4685 | 0 |
| *POS_-4_A* | 0.8874 | 0.1377 | 6.4422 | 0 |
| *Free Energy* | 0.084 | 0.0166 | 5.0552 | 0 |
| *POS_-3_G* | −0.7325 | 0.1754 | −4.1762 | 0 |
| *POS_5_G* | −0.6234 | 0.1555 | −4.0078 | 1e−04 |
| *POS_3_C* | −0.6688 | 0.1757 | −3.8056 | 1e−04 |
| *POS_3_A* | −0.3413 | 0.1189 | −2.8703 | 0.0041 |
| *GCcontent* | −2.5733 | 0.9705 | −2.6515 | 0.008 |
| *POS_6_C* | −0.3357 | 0.1639 | −2.0478 | 0.0406 |
| *POS_-4_C* | −0.4127 | 0.2031 | −2.0325 | 0.0421 |
| *POS_-3_C* | −0.3276 | 0.1852 | −1.7684 | 0.077 |
| *POS_6_A* | 0.2212 | 0.1271 | 1.7402 | 0.0818 |
| *POS_5_A* | −0.2167 | 0.1317 | −1.6456 | 0.0999 |
| *POS_-4_G* | −0.219 | 0.1806 | −1.2127 | 0.2253 |
| *POS_5_C* | −0.1851 | 0.1591 | −1.1635 | 0.2446 |
| *POS_6_G* | −0.1877 | 0.1665 | −1.1274 | 0.2596 |
| *POS_-3_A* | −0.1183 | 0.1251 | −0.9461 | 0.3441 |
| *(Intercept)* | 0.3527 | 0.4083 | 0.8638 | 0.3877 |

**Table S2.**
Summary of coefficient estimates for variables in the logistic regression model.

**Table S3**
MASTER-seq quantifications and model fitted values across ACA sites in yeast.

**Table S4**
MASTER-seq quantifications in mESCs and in EBs across sites in mouse.

**Table S5**
List of strain genotypes.

**Table S6**
List of sequences used throughout the experimental protocols present in the work.
